# Pathways of savannization in a mesic African savanna-forest mosaic following an extreme fire

**DOI:** 10.1101/2021.05.27.445949

**Authors:** Heath Beckett, A. Carla Staver, Tristan Charles-Dominique, William J. Bond

## Abstract

Fires in savannas limit tree cover, thereby promoting flammable grass accumulation and fuelling further frequent fires. Meanwhile, forests and thickets form dense canopies that reduce C4-grass fuel loads and creating a humid microclimate, thereby excluding fires under typical climatic conditions.
However, extreme fires occasionally burn into these closed-canopy systems. Although these rare fires cause substantial tree mortality and can make repeat fires more likely, the long-term consequences of an extreme fire for closed canopy vegetation structure and potential to convert to savanna (hereafter “savannization”) remain largely unknown.
Here, we analysed whether an extreme fire could, alone, alter species composition, vegetation structure, and fire regimes of closed-canopy ecosystems in an intact savanna-forest-thicket mosaic, or whether successive fires after an initial extreme fire were necessary to trigger a biome transition between from forest to savanna.
We found that forests that only burned once recovered, whereas those that burned again following an initial extreme fire transitioned from closed-canopy forests towards open, grassy savannas.
While thickets had less tree mortality in fires than forests, repeat fires nonetheless precipitated a transition towards savannas.
Colonization of the savanna tree community lagged behind the grass community, but also began to transition.

**Synthesis:** Our results suggest that rare extreme fires, followed by repeated burning can indeed result in savannization in places where savanna and forest represent alternative stable states.

## Introduction

Fires are becoming more frequent and severe in forested ecosystems globally (Andela *et al.*, 2019, Jolly *et al.*, 2015). In tropical forests, extreme fires associated with droughts and agriculture are expanding into forests that historically burnt only once every few hundred years (Chen *et al.*, 2014, Morton *et al.*, 2013, Sanford *et al,* 1985). Some authors have hypothesized that these tropical forest understory fires may result in runaway feedbacks, leading to more frequent fires and to savannization of forests, resulting in eventual forest collapse (Cochrane, 1999, Barlow & Peres, 2004, Barlow & Peres, 2008, Silverio et al, 2013, Flores *et al*, 2016, van Nes et al, 2018). However, although we know that the occurrence of one forest understory fire can increase the likelihood of subsequent fires, direct evidence of forest savannization by runaway feedbacks is sparse. Invasive exotic grasses have been shown to invade the forest understory only locally (Silverio *et al.*, 2013), limited by dispersal (although accelerated by human activity (see Veldman and Putz, 2010)) and possible biogeochemical feedbacks. As a result, the question of whether fires alone can convert an intact tropical forest to a savanna remains open.

Historically and today, fires burn frequently in most savannas, at a rate of several per decade, but rarely penetrate beyond intact forest margins (Kellman & Meave, 1997, Biddulph & Kellman, 1998, Hennenberg *et al.*, 2008). In some places, this can allow the persistence of a forest-savanna mosaic, despite the potentially destructive impact of fire on forest trees (Cochrane & Laurance, 2002, Brando *et al,* 2011, Brando *et al,* 2014), by feedbacks that are spatially localized (Favier *et al.*, 2004, Beckett & Bond, 2019). Mechanistically, C4-grass-fuelled fires maintain an open canopy by preventing fire-sensitive forest saplings from establishing, but fires stop readily at savanna-forest boundaries, because shade in the forest understory prevents shade-intolerant C4 grasses from accumulating within meters of the boundary (Charles-Dominique *et al.*, 2018, Hennenberg *et al.*, 2008, Hoffmann *et al.*, 2012a). Forest microclimates also reduce wind speeds and increase fuel moisture (Biddulph & Kellman, 1998, Hoffmann *et al.*, 2012b, Little *et al.*, 2012), further hampering fire spread. These feedbacks provide support for the idea that savanna and forest are alternate stable states (Staver *et al.*, 2011a). However, the impact of extreme events (especially extreme fires, see Keeley & Pausas, 2019) on the dynamics of these mosaics is largely unknown.

One possibility is that mosaics are mostly stable, with distinct distributions and stable edges, not just on short to medium time scales (ten to several hundred years), but also on longer time scales (Killeen *et al*, 2006, Breman et al, 2012, Rull *et al*, 2013, Cardoso *et al*, 2020). This might be the case if savanna-fire feedbacks are very strong (Beckage et al, 2009, Dantas et al, 2016), or if some other process besides fire – e.g., oligotrophy, seasonal flooding – play a strong role in stabilizing savanna-forest distributions within a mosaic (Veenendaal et al, 2015, Murphy and Bowman, 2012). If this is the case, extreme fires in forests may have minor medium-term effects on forest structure, acting as a stand-replacing disturbance from which forests recover via succession (Fearnside, 1990, Kennard, 2002, Brando et al, 2020) rather than as a trigger of a catastrophic transition.

An alternative possibility is that savanna-forest boundaries may be highly dynamic over longer time-scales (Schwartz *et al.*, 1986, West *et al.*, 2000, Staver *et al*, 2017, Aleman et al 2019) and in response to changing climate or climate events (Hirota *el al,* 2010, Breman et al, 2012, Beckett & Bond, 2019, Wright et al, 2021, Sato *et al,* 2021). Spatially explicit modelling work suggests that fires at the savanna-forest boundary can influence the biogeographic distribution of biomes (Staver *et al.*, 2011b, van de Leemput *et al*, 2015, Goel *et al,* 2020a, Goel *et al,* 2020b, Wuyts *et al,* 2019). Meanwhile, local observations confirm that boundaries can be dynamic through time (Brook & Bowman, 2006, Silva *et al*, 2008, Ibanez *et al*, 2013, Beckett & Bond, 2019), from Africa (Baccini et al., 2017; Aleman et al., 2018), to South America, to Australia (Marimon et al., 2014; Ondei et al., 2017; Stevens et al., 2017; Rosan et al., 2019).

Modern evaluations of invasions of forests by savannas have predominantly focused on the Neotropics (Cavelier et al, 1998, Veldman and Putz, 2011, Silverio *et al.*, 2013, Flores et al 2016), typically in the context of invasive exotic grasses, El Niño-fuelled droughts, and deforestation (Nepstad *et* al. 1996, Barlow and Peres, 2004, Barlow and Peres, 2008, Veldman and Putz, 2011, Silverio *et al.*, 2013). This work provides invaluable insights into the potential processes of savannization, which includes the susceptibility of forest trees to fire induced mortality (Barlow and Peres, 2008, Staver *et al*, 2020), the invasion of grassy fuels following repeat fires (Veldman and Putz, 2011, Silverio *et al*, 2013), and finally the formation of no-analogue grassy systems that differ from old-growth savannas (Veldman and Putz, 2011). However, these transitions involve exotic grass species, which lead to novel fire regimes, and deforestation-derived boundaries (Alencar *et al.* 2015), with ambiguous historical and palaeoecological precedents.

From its conception in the 1940’s (Aubreville, 1949, Budowski, 1956), the term savannization has been used to refer to fire-induced degradation of tropical forests via anthropogenic deforestation and agriculture (Nyerges, 1990, Uhl *et al*, 1982, Cochrane, 1999, Nepstad *et al*, 1999). The term re-emerged following predictions of large-scale Amazonian Forest collapse under elevated CO_2_ (Cox *et al*, 2004) and is widely used in the contemporary literature to refer to a reduction of forest tree biomass and species. In doing so, forest ecologists – unintentionally but incorrectly – have maligned savannas as degraded, species-poor forests or as an early successional stage of forest vegetation (Veldman, 2016, Bond, 2016, Schmidt *et al*, 2019). The term savannization should be used to refer to the conversion of one vegetation state (typically forest) into a savanna, a system subject to the ecological processes that determine savanna vegetation dynamics. This replacement of forest with savanna (true ‘savannization’) could, in theory, be possible in regions where savannas and forests represent alternative stable states (Staver & Levin, 2012, Archibald *et al*, 2019, Aleman *et al*, 2020). However, the question of whether extreme fires can cause a biome switch from intact forests to old-growth savannas has not yet been settled.

This question has both theoretical and practical importance. From the perspective of understanding ecological processes, we have no clear understanding of the possible role of extreme fires in biogeographic transitions between savanna and forest *(e.g.,* Aleman et al 2019) and in shaping transitions in response to changing climate (Beckett & Bond 2019). From a practical perspective, recent decades have seen significant increases in deforestation and fragmentation of intact tropical forests (Watson et al, 2018; Taubert et al, 2018). The most commonly proposed solution to this is to plant trees in order to restore fragmented landscapes (e.g., the Bonn Challenge, REDD+; Veldman et al, 2015b), but this can be harmful to the natural, biodiverse, open systems found in savanna-forest mosaics (Bond and Parr, 2010, Veldman et al, 2015b, Ratnam et al, 2011). Understanding the stability and dynamics of mosaics would help to justify and inform their management as mosaics, with a focus on the preservation of savannas as well as forests (Bond & Parr, 2010).

Here, we have examined vegetation trajectories after extreme fires empirically, focusing on a mesic savanna-forest-thicket mosaic in Hluhluwe iMfolozi Park, South Africa. Extreme fires burned into forests and thickets in 2004 and 2008, some of which have burned again subsequently and others of which have not. These events offer a natural experiment in which to observe, *in situ,* the ecological trajectories of closed-canopy vegetation following extreme fires. On the one hand, a forest or thicket experiencing fire might recover its original state. If so, we expect 1) a reduction in tree cover and biomass that is only temporary, 2) the inhibition of a regular fire regime (e.g. by suppression of grassy fuels), and 3) a transient shift to early-successional forest trees species (light-loving but not as fire tolerant as savanna species) that eventually transitions back to the original forest state. On the other hand, a forest or thicket might transition to savanna. If so, we instead expect 1) a reduction in tree cover and biomass that persists through time, 2) the establishment of a regular fire regime (or at least of sufficient grassy biomass to support one), and 3) a shift in tree species composition, from shade-tolerant but fire-sensitive species that compete for light, to shade-intolerant species that invest in fire protection. Differentiating between these possibilities provides us the rare opportunity to test the forest savannization hypothesis developed for the Amazon (Cochrane et al 1999), allowing us to investigate what happens in the aftermath of extreme disturbances in forests and thickets.

## Materials and Methods

The northern section of Hluhluwe-iMfolozi Park (HiP) receives sufficient annual rainfall (approx. 1000mm per annum) (Balfour & Howison, 2002) to support a closed-canopy forest, with vegetation dynamics that are driven by fire compared to the more arid (approx. 600mm per annum), herbivore-driven system in the iMfolozi section (Staver *et al.*, 2012). Closed-canopy vegetation types in HiP are differentiated into scarp forests (which we refer to as forests here onwards) and thicket (*sensu* Charles-Dominique et al 2015a). Forest, thicket, and savanna definitions are based on the classification scheme used in Charles-Dominique et al (2015a), using vegetation structure and associated growth forms (Woodward et al, 2004; Ratnam et al, 2011). In HiP, forests are classified as tall (>10m) woody vegetation with a shade-tolerant intermediate tree layer, lacking a C4 grass layer but occasionally containing patches of C3 grasses and herbaceous plants among the litter layer (Charles-Dominique *et al*, 2015). Thickets are instead classified as having dense shrub and treelet vegetation, a canopy that is generally 4 to 6m tall, and a variable understory with dense patches of herbaceous sub-shrubs, shrubs and occasional patches of C4 grass subtypes that are shade-tolerant (Charles-Dominique *et al*, 2015, Charles-Dominique *et al*, 2018); thicket in the HiP context is entirely distinct from and not to be confused with subtropical Albany thicket, whose distribution is centered in the Eastern Cape of South Africa. Finally, savannas are classified as having discontinuous tree cover with a continuous layer of C4 grasses. Forest and thicket patches coexist with savanna patches in a mosaicked landscape, which is home to a full complement of savanna herbivores, including elephant and black rhino.

As in most of southern Africa, fires are usually lit by people, pre-empting lightning fires which generally occur near the end of the dry season (Archibald *et al.*, 2017). Humans have been burning fires in the region for at least 150 ka with Iron Age farmers arriving in the area ~ 2 ka (Hall, 1980, Staver *et al.*, 2017). The Hluhluwe section of HiP experiences relatively frequent fires, with mean and median fire return intervals of 2.9 and 1.3 years, respectively (Balfour and Midgley, 2008).

On the 14^th^ and 15^th^ of September 2008, an extreme fire occurred in the Hluhluwe section of HiP that, unlike typical savanna fires, burned into large tracts of thicket and forest. These extreme fires resulted from a combination of air temperatures above 30 °C, relative humidity below 30%, and wind speeds averaging more than 25 km.hr^-1^ with gusts to 30 km.h^-1^ or more (Bradstock et al 2009; Browne & Bond, 2011). In this region, such conditions are exceedingly rare: over the period 2001-2008, a total of only 69.6 hours (~ 3 days) of extreme fire weather conditions occurred, with the longest consecutive period (11 hours) occurring on the day of the extreme fire in 2008 (Browne & Bond, 2011). Aerial photography of the region from 1937 to 2013 shows that, from 1937 to 1992, closed canopy patch sizes gradually increased in extent (Beckett & Bond, 2019). However, between 1992 to 2013, when this extreme fire occurred, forest and thicket patches were lost from the landscape. Some areas that experienced this extreme fire burned subsequently, whereas others did not (see below for further detail, Appendix S1).

### Vegetation sampling

We used a set of plots in which the structure and species composition of woody plants were recorded in forest, thicket and savanna vegetation types. A total of 131 plots were sampled (Fig. 1), 51 of which were first sampled in 2007 by Staver *et al* (2012) and resampled in 2014 by Case and Staver (2017), 31 of which were sampled once in 2014 by Charles-Dominique *et al* (2015a), and an additional 49 which were established specifically for this study between 2009 and 2015. Staver et al (2012) sampled 253 plots across HiP, 51 of which were chosen as the savanna baselines as they burnt in the 2008 fire and were near forest and thicket patches. Intact forest and thicket plots were established and sampled in 2014 and 2015 in mature unburnt vegetation patches (Fig. 1), as mapped by Whateley and Porter (1983) and by aerial-photograph-derived vegetation maps (Beckett & Bond, 2019). These represent our pre-fire baselines for forest and thicket communities and, along with the 2007 savanna plots, are referred to from here on as our ‘before’ dataset, while burnt plots represent the ‘after’ dataset.

**Figure 1.**
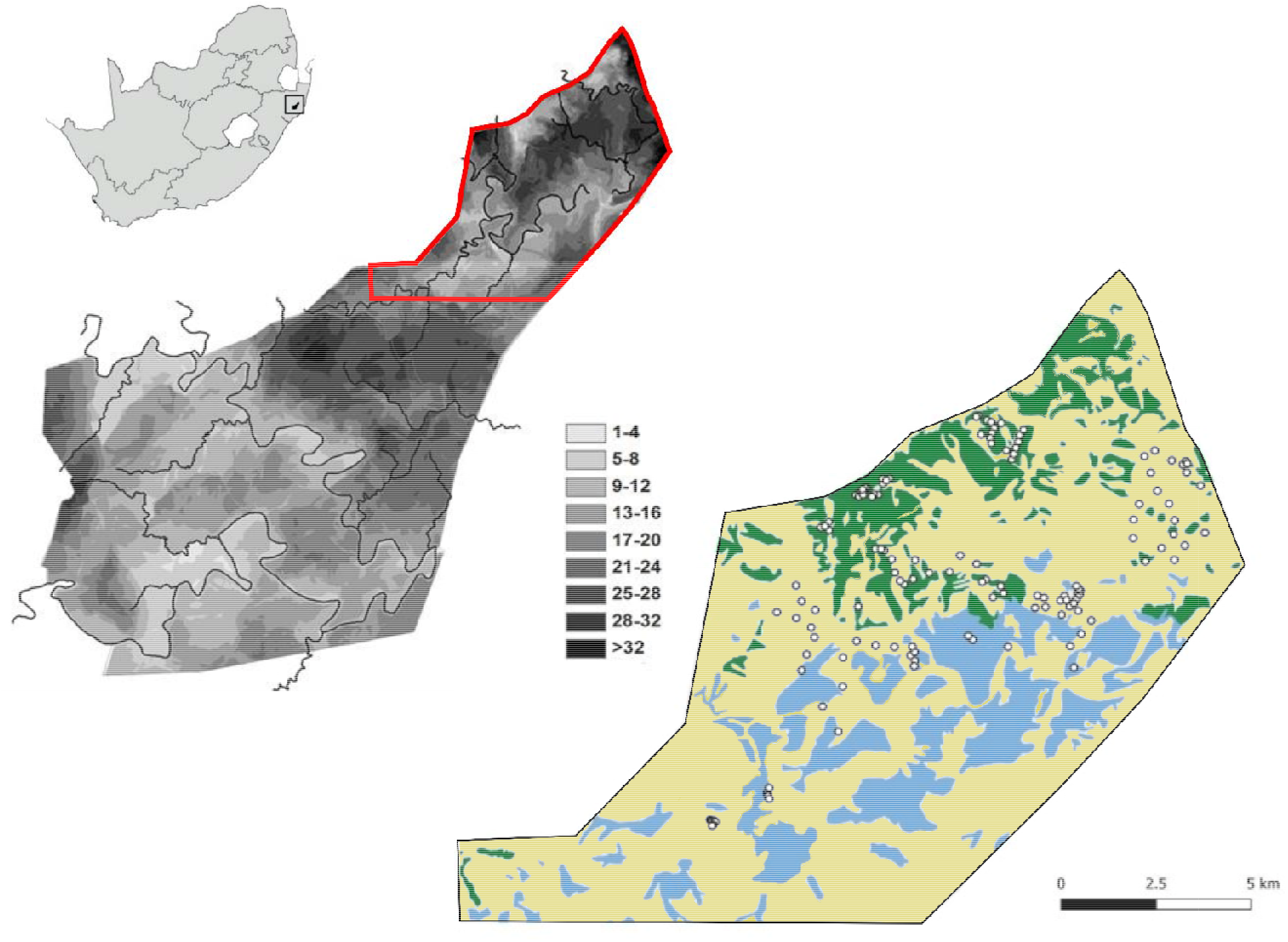
Fire Frequency map (left) of Hluhluwe iMfolozi Park from 1955 to 2013 based on Hluhluwe iMfolozi Park records with the Hluhluwe section outlined in red, and (right) the distribution of plots within the Hluhluwe portion of HiP and biomes based on a reclassification of Whateley & Porter (1983) by Charles-Dominique *et al* (2015). Green represents forest, blue represents thickets, and yellow represents savannas. Inset map shows the location of Hluhluwe iMfolozi Park within South Africa.

To determine vegetation change trajectories after the extreme fires, in 2013, we identified regions within the 2008 fire scar that were known to have burnt subsequently, and established plots in these areas. These plots were sampled repeatedly, supplemented with some additional, once-sampled plots allowing a space-for-time chronosequence to evaluate vegetation change. Three additional plots were laid out in a forest that burned in a separate fire in 2004 and sampled nine and 11 years after the fire. These three plots (with 20 m buffers on all sides) covered approximately 80 % of the burnt forest area. One of these plots was burnt again in 2012 and was excluded from the 11-year resampling as it had no analogues. The data from nine years after the fire are included in the analyses, while the repeat sample at 11 years are only included in figures in the Supplementary Materials for interest. Plot sampling dates, repeat sample dates, and vegetation history are summarized in Appendix S1.

Each plot measured 40 m x 10 m, within which area we identified every woody plant (including lianas) above 50cm in height and recorded its height as one of four size classes (0.5-2 m, 2-5 m, 5-10 m, and >10 m). In 103 of the 134 sites, the grass layer was evaluated at 20 points along the plot midline; grass biomass was measured via disk pasture meter (DPM) and the dominant grass species beneath the disk was recorded. A DPM is a weighted metal disk of a known diameter, which is dropped on a grass sward from a standard height. The resting height is a measure of the grass biomass under the disk (Bransby and Tainton, 1977), which can be calibrated locally (grass biomass = 1.26 + 26.l*[DPM], R2 = 0.73, N = 1745; Waldram et al, 2007).

### Fire occurrence

Fire frequency of each plot from 2001 to 2016 was derived from a combination of the Ezemvelo KwaZulu Natal Wildlife fire records for HiP and personal observations. The EKZNW fire records are hand drawn maps from rangers responsible for sections of the park. These indicate the general location of the fire, but do not include information on fine-scale fire refugia and unburnt vegetation patches within the burned area (see Fig. 1). Where EKZNW fire records indicated fires burned into large forest patches, we verified these records against MODIS Active Fire Detections (MCD14DL, Collection 6). MODIS Active Fires maps thermal anomalies within 1km pixels using the MODIS MOD14/MYD14 Fire and Thermal Anomalies algorithm (Giglio *et al.*, 2003). MODIS Active Fires increases the probability that fires within forests are detected, since forest canopies can interfere with detection of burn scars.

### Data analysis and statistics

Aerial photography and vegetation maps were analysed using Quantum GIS 3.0.0 (Quantum GIS Development Team, 2018). Statistical analyses were performed in R (version 3.4.0, R. Core Team, 2017), using the packages *vegan* (Oksanen *et al.*, 2013) and *tidyverse* (Wickham, 2017). Fire frequency before and after extreme fire was compared using a Wilcoxon signed-rank test for non-parametric paired data. Grass biomass was compared between vegetation types and time periods using a one-way analysis of variance with Tukey’s HSD contrasts to identify significant differences in mean values. We used a multivariate analysis of variance (MANOVA) to test for differences in the total mean cumulative length of the vegetation types at different time periods and followed this with a series of ANOVA models to test for significant differences between vegetation types within the size classes. Cumulative length is a composite measure of species importance, similar to basal area, which combines stem density (reported in Appendix S2) and tree height into a single value (Fei *et al.*, 2005). This is done by summing the heights of all individuals of a species, thereby allowing large trees to contribute more to the overall score than small trees. We performed an NMDS Ordination on a Bray-Curtis species dissimilarity matrix of the cumulative length of each tree species for the ‘before’ plots, representing vegetation communities that had not experienced the extreme fire (the intact forest, intact thicket, and 2007 savanna plots). Data were transformed using a Hellinger transformation (Legendre & Legendre, 2012). Species were assigned to a biome (forest, thicket, or savanna) based on their association with sites in the NMDS ordination; species not found before the extreme fire were classified as pioneers. We then performed a Principal Coordinates Analysis (PCoA) and a pairwise permutational MANOVA on a Bray-Curtis dissimilarity matrix of the total cumulative length in each plot, as well as within the size classes to identify changes in the composition of species types.

## Results

### Establishment of a savanna fire regime

Forests and thickets rarely burned, with intact forest and thicket sites experiencing no fires between 2000 and 2016 (Fig. 2). Savanna sites, in contrast, burned several times in a decade. In fact, fire frequency (FF) increased from approximately one fire every four years (FF = 0.24 yr^-1^ (SD = 0.11)) before the 2008 extreme fire to one fire every three years (FF = 0.36 yr^-1^ (SD = 0.13); t = −11.8, df = 43, p < 0.05) after 2008. Thicket patches that burned in the 2008 extreme fire quickly established a savanna fire regime, with sites burning on average once every three years (FF = 0.34 yr^-1^ (SD = 0.07)), up from no fires between 2000 and 2007.

**Figure 2.**
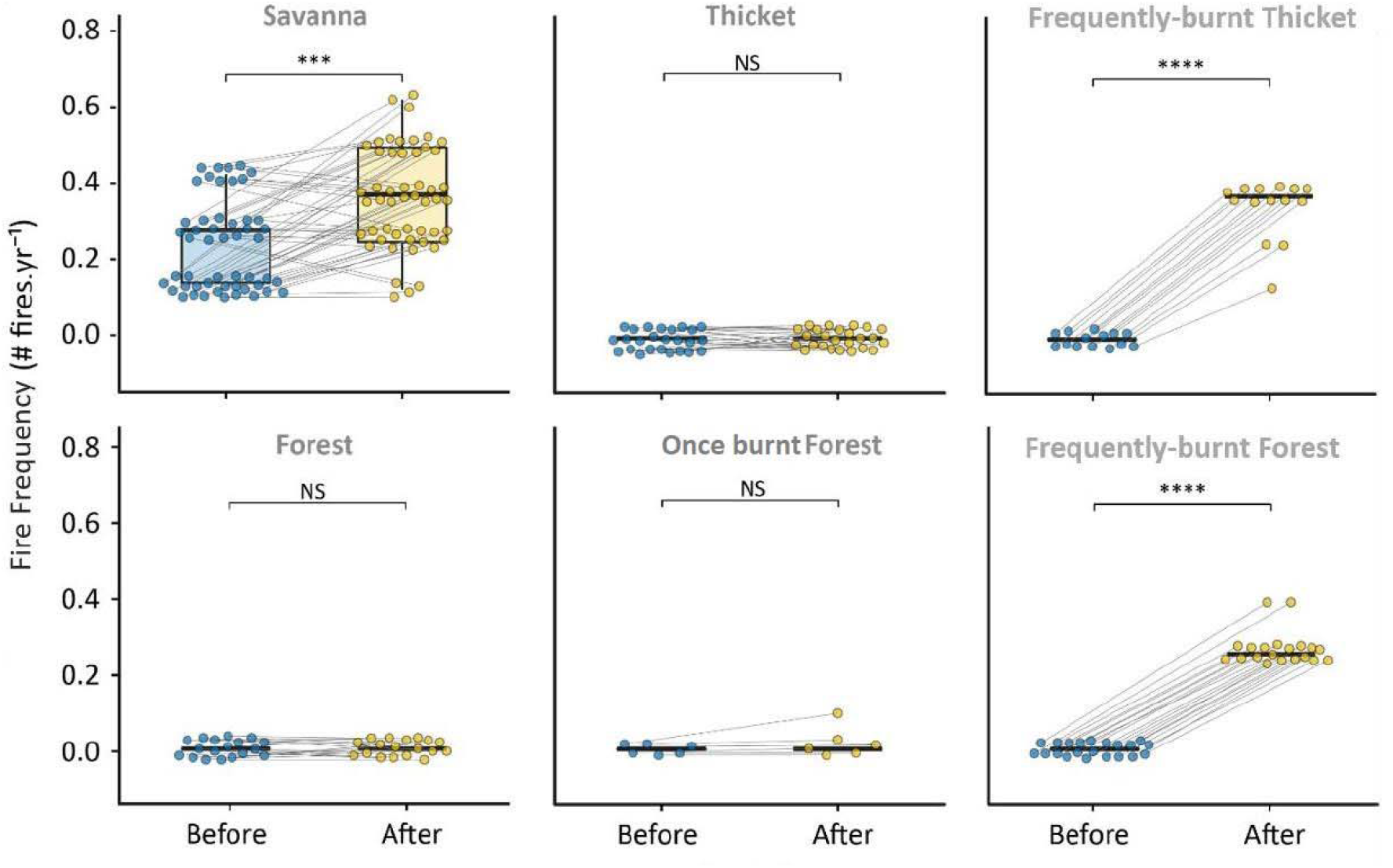
Fire Frequency for plots within each vegetation type (indicated by grey text) before and after extreme fires (not including the year of the extreme fire), calculated using the MODIS Active Fires Product from 2000 to 2016. Grey lines link plot points before and after extreme fires. Asterisks indicate significance level (*** <0.05, **** < 0.001, NS = no significant difference).

Some forests that burned in extreme fires burned again, whereas others did not. To investigate how contrasting fire returns affected their post-extreme-fire trajectories, we split plots into two groups, those that burned only once (hereafter “once-burnt forests”) versus those that burned in an extreme fire and at least once subsequently (hereafter “frequently-burnt forests”). Frequently-burnt forests burned on average once every four years after the extreme fire (FF = 0.26 yr^-1^ (SD = 0.04), Fig. 2).

### Colonisation by savanna grasses

Intact thicket and forest plots differed from savanna plots in having closed canopies and little or no grass in the understory (Fig. 3). Savanna plots had on average 233 gm^-2^ (SD = 87) of grass biomass whereas intact thicket and forest patches had 148 gm^-2^ (SD = 84) and 10 gm^-2^ (SD = 2.8) respectively (Fig. 3). Frequently-burnt thicket and frequently-burnt forest plots showed an increase in grass biomass following successive fires, with grass biomass four and 22 times higher than intact thicket and forest plots respectively (Fig. 3). Once-burnt forest plots had relatively high, but variable, grass biomass (130 gm^-2^ (SD = 114)) in the first year after the extreme fires. By the end of the study, 9 years later, grass biomass was less variable (33 gm^-2^ (SD = 11)), approaching the levels of intact forest grass biomass. In terms of grass species, *Panicum maximum* colonised forests and thickets immediately after fire, whereas *Themeda triandra* increased in abundance only where fires occurred repeatedly (Fig. 3).

**Figure 3.**
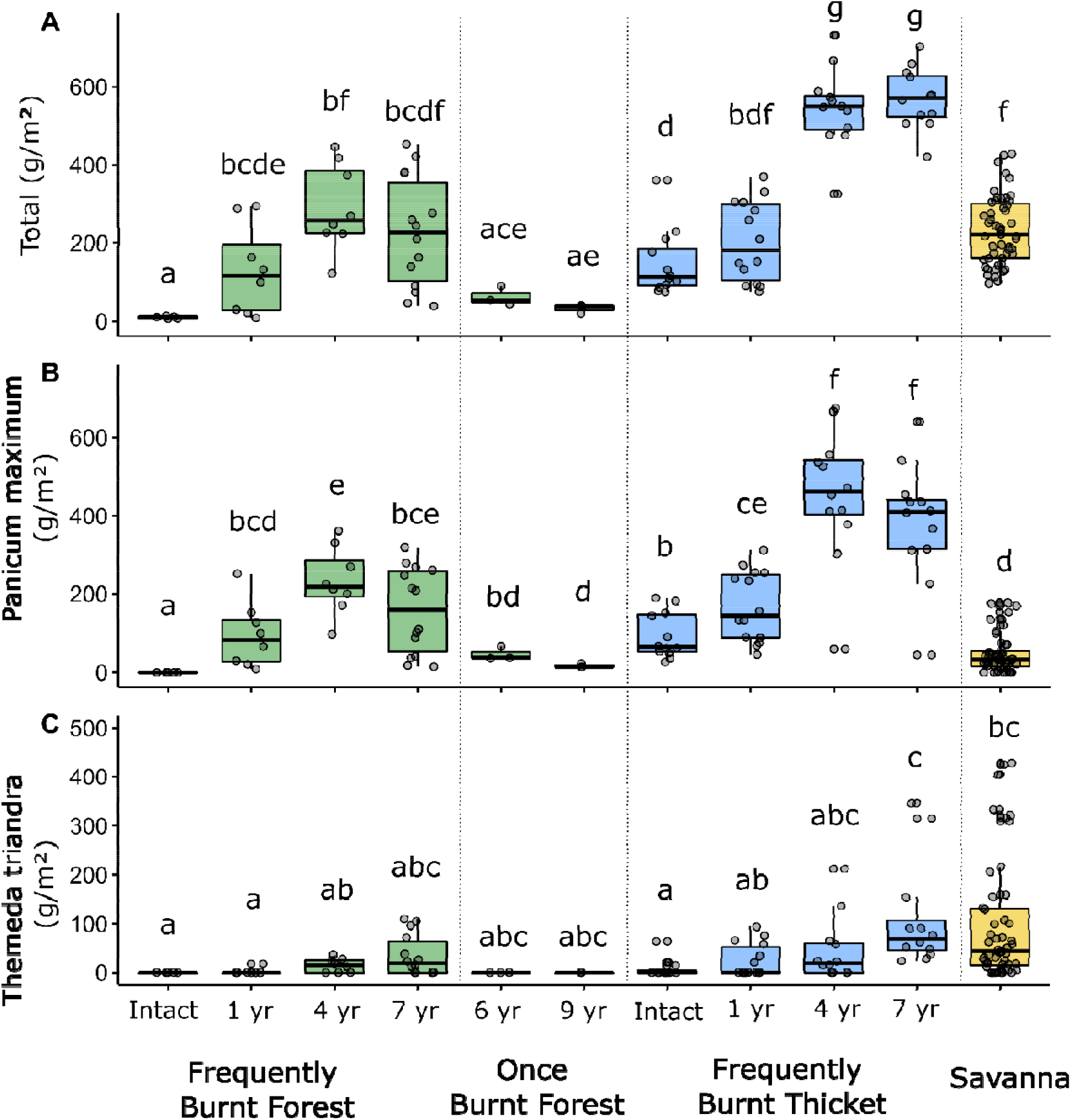
Grass biomass in frequently-burnt forests, once-burnt forest, frequently-burnt thicket and savanna, (a) in total, (B) of *Panicum maximum,* and (C) of *Themeda triandra.* Green bars represent forest plots, blue represents thicket, and yellow represents savanna. Error bars indicate standard errors. Points (plot grass biomass) are included to display variation. Shared letters indicate lack of a significant difference within panels based on Wilcoxon signed-rank test, p < 0.05.

### Trajectories in woody structure

Forest, thicket, and savanna plots displayed distinct woody size-class distributions (See Appendix S3), with most savanna trees in the smallest size class, most forest trees in the largest size class, and an even spread between the two intermediate size classes in thicket (Fig. 4, Appendix S3). Savanna plots showed little change in structure between 2007 and 2014, despite an increase in fire frequency (Pillai’s Trace = 0.039, approx F(3,91) = 1.24, p = 0.301, Appendix S3).

**Figure 4.**
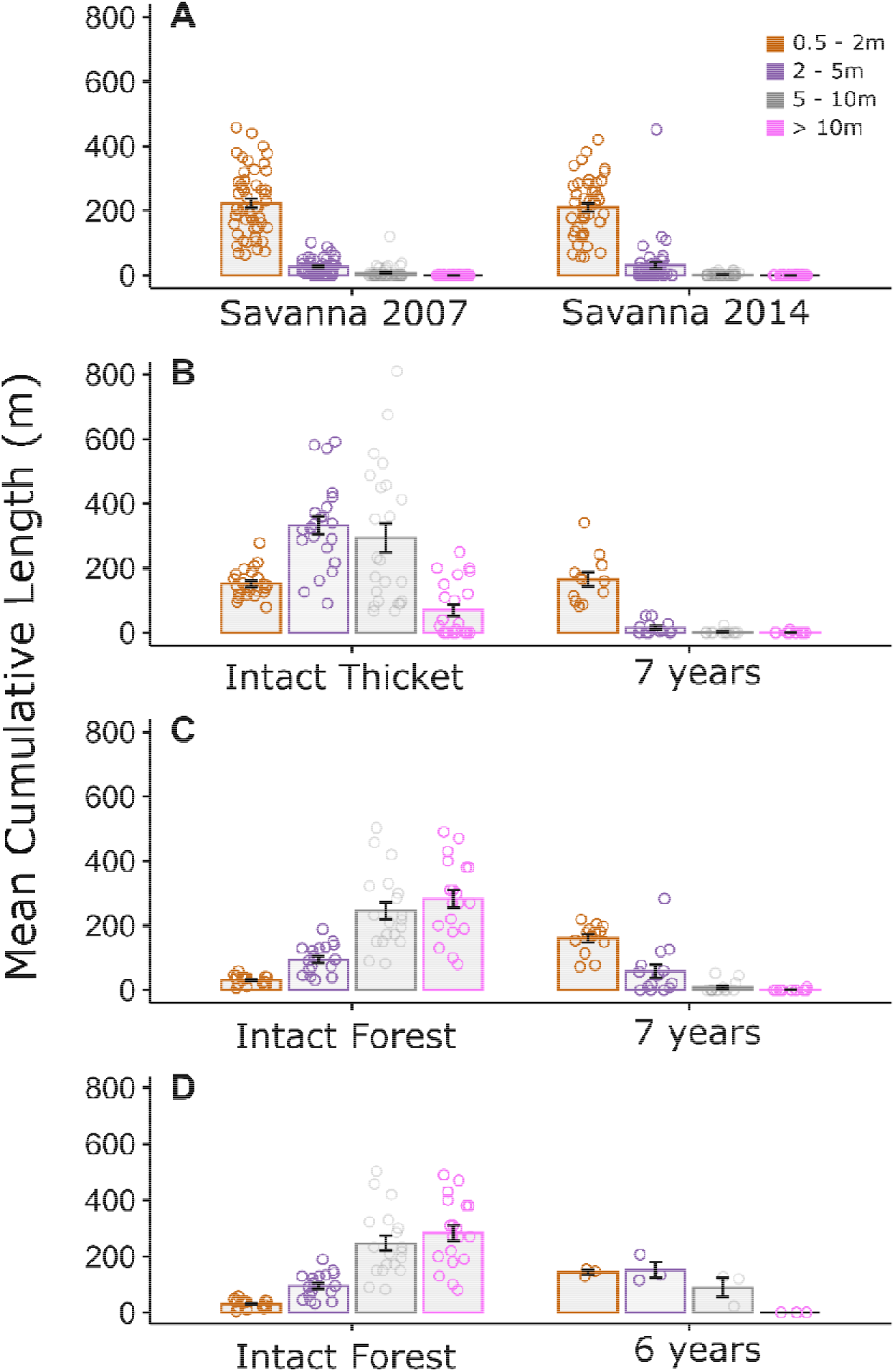
Mean cumulative length of all tree species within the four size classes ‘before’ and ‘after’ the extreme fire in A) savanna, B) frequently-burnt thicket, C) frequently-burnt forest, and D) once-burnt forest. Error bars indicate standard errors. Points represent individual plots.

Thicket plots that burned frequently were structurally distinct from intact thicket plots (Pillai’s Trace = 0.76, approx F(4,30) = 23.41, p < 0.001, Appendix S3), converging to savanna woody size class structure 7 years after the extreme fire (Pillai’s Trace = 0.15, approx F(4,51) = 2.24, p = 0.077, Appendix S3), with most trees in the smallest size class. Similarly, forests that burned multiple times were different to intact forests (Pillai’s Trace = 0.89, approx F(4,28) = 58.73, p < 0.001) and also converged to a savanna-like vegetation structure (Pillai’s Trace = 0.20, approx F(4,53) = 3.37, p = 0.016) 7 years after the extreme fire, with comparable numbers of trees in the smaller two size classes but differences in the larger two size classes (Appendix S3).

Six years after the 2008 extreme fire, once-burnt forests were structurally distinct from both intact forests (Pillai’s Trace = 0.902, approx F(4,17) = 38.90, p < 0.001) and savannas (Pillai’s Trace = 0.77, approx F(3,43) = 46.72, p < 0.001). The six year once-burnt and the frequently-burnt forests were also different (Pillai’s Trace = 0.61, approx F(4,12) = 4.72, p = 0.016), with the second largest size class contributing to the majority of the difference between the two (F(1,15) = 21.96, p < 0.001). Nine years after an extreme fire, statistical differences between once-burnt forests and intact forests (Pillai’s Trace = 0.761, approx F(4,17) = 13.50, p < 0.001), savannas (Pillai’s Trace = 0.767, approx F(3,43) = 47.30, p < 0.001), and frequently-burnt forests (Pillai’s Trace = 0.716, approx F(4,12) = 7.57, p < 0.05) were qualitatively similar to those of the six-year once-burnt forests (Appendix S3, Appendix Fig. S1), except that the cumulative length of the second largest size class of once-burnt forests began to approach that of the intact forests (F(1,20) = 3.64, p = 0.071).

### Trajectories in woody species composition

Forest, thicket, and savanna plots were characterized by distinct woody species assemblages (Fig. 5, Appendix Table S4 & Table S5). The overall (Appendix Table S4) and size class specific (Appendix Table S5) abundance of species types within savannas did not differ between 2007 and 2014 (Fig. 6).

**Figure 5.**
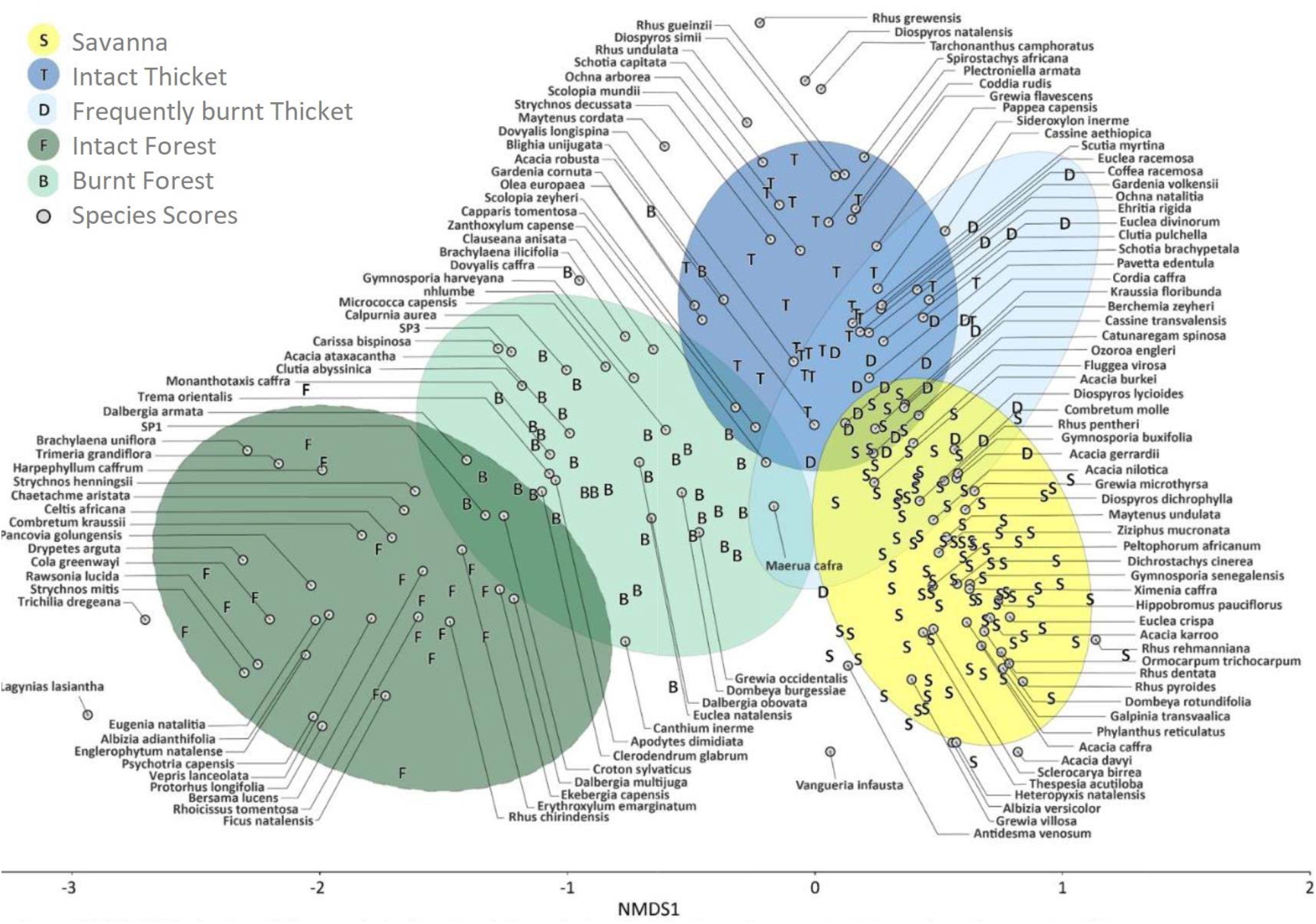
NMDS ordination of the cumulative length of all species in all height classes in plots. Ellipses show the grouping of plots. Species associations used in the composition analyses are based on species found in plots before the extreme fires only. Letters indicate plots, and grey dots indicate species.

**Figure 6.**
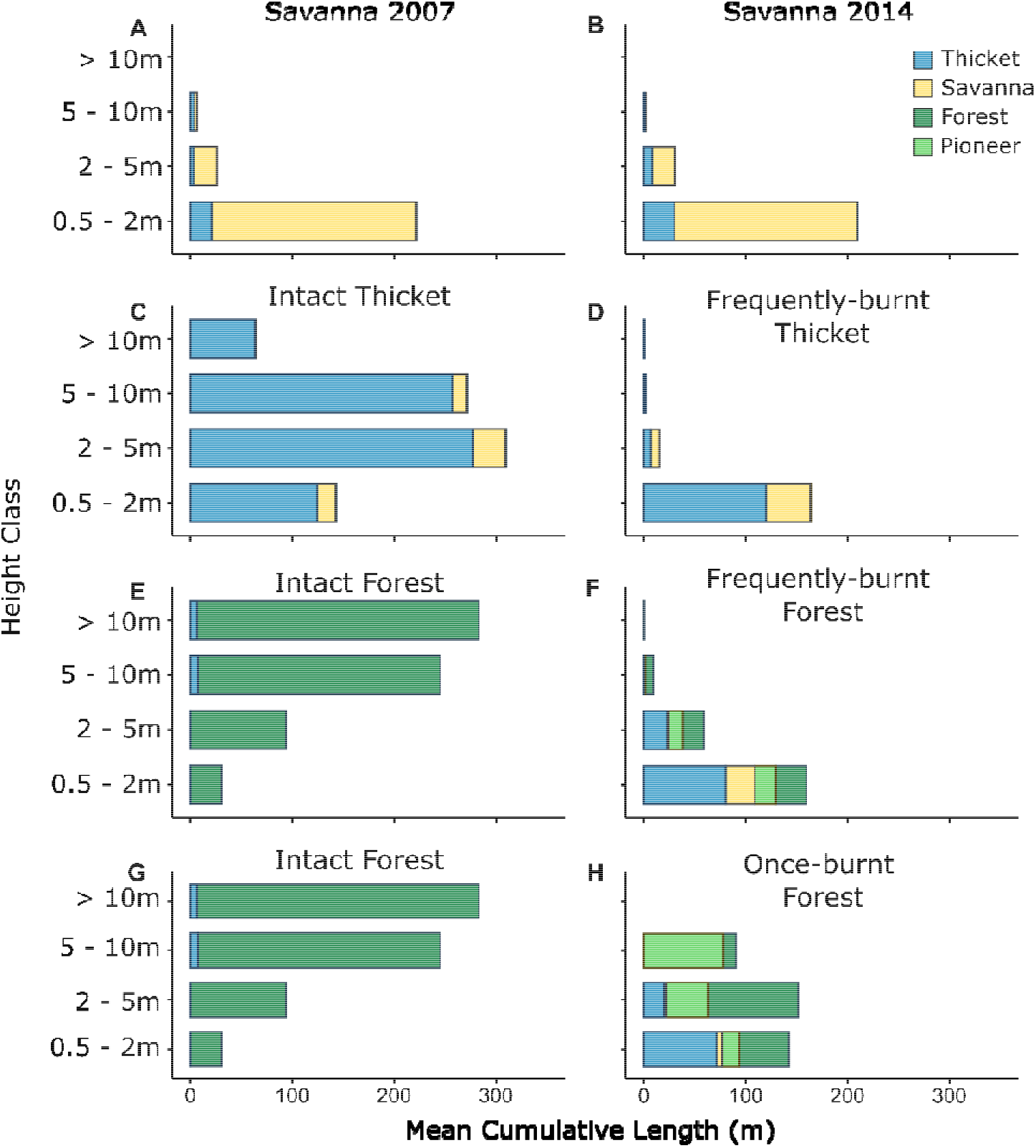
Initial and final composition and structure of savannas in 2007 (A) and 2014 (B), intact thicket (C) and frequently-burnt thicket 7 years after extreme fire (D), intact forest (E) and frequently-burnt forest 7 years after extreme fire (F) and intact forest (G) and once-burnt forest 6 years after fire (H). Panels display the average cumulative length of tree species within each size class for the different time periods ‘before’ and ‘after’ the extreme fire. Colours indicate the biome in which these species are predominantly found. Pioneer species are those which were not found in plots sampled before the extreme fire. Data recorded in intact forest and thicket vegetation represent pre-fire space for time substitutions.

Compositionally, frequently-burnt thickets differed from savannas 7 years after the extreme fire (Fig. 6, Appendix Table S4 & S5), despite both vegetation types experiencing similar disturbance regimes. This was most evident in the smallest size class, in which frequently-burnt thickets were dominated by basally resprouting thicket species, *e.g., Euclea racemosa, Euclea divinorum* and *Berchemia zeyheri,* mixed in with some savanna species, including *Acacia karroo (=Vachellia karoo), Acacia nilotica (=V. nilotica), Dichrostachys cinerea, Cordia caffra, Combretum molle* and *Rhus (= Searsia) pentheri* (Fig. 5), whereas savannas rarely featured thicket species.

By contrast, while frequently-burnt forests also differed from savannas (Appendix Table S4 & S5) 7 years after the extreme fire, their composition had changed more dramatically. Forests, when frequently burned, completely switched to being dominated by thicket species, including *Euclea natalensis* and *Euclea racemosa,* and savanna species (see Fig. 5 & 6), and closely resembled the frequently-burnt thickets.

Once-burnt forests were dominated by pioneer forest species seldom found in intact forests (including *Trema orientalis, Croton sylvaticus,* and *Dombeya burgessiae).* These trees apparently grew from seed banks, which contrasts with trees in frequently-burnt thicket plots, which may have resprouted basally. After 6 years, these once-burnt forests recovered some species characteristic of mature forest, including *Englerophytum natalense* and *Chaetacme aristata,* in smaller size classes, although the larger size classes remained dominated by the unique pioneer forest assemblage (Fig. 6). The once-burnt forests (after six years) were not different to the frequently-burnt forests in overall abundance of species types (Appendix Table S4), but they were different when considering size-class-specific species-type abundances (Appendix Table S5). Nine years after fire, once-burnt forests were dominated by mature forest species in the two smaller size classes and by pioneer forest species in the second largest size class (Appendix Fig. S2). These plots differed from frequently-burnt forests in overall and size-class-specific species-type abundances (Appendix Table S4 & S5, Fig. S3.).

## Discussion

Extreme fires differ from typical savanna fires in penetrating beyond the biome boundaries, often opening up the canopy of closed forest or thicket. Here, we found that forests that burned only once recovered via secondary successional pathways, whereas forests and thickets that burned again following the initial extreme fire underwent a persistent transition from closed-canopy vegetation states to open-canopy grassy systems. Among grasses, *Panicum maximum* invaded first but was gradually replaced by *Themeda triandra,* a more flammable species (Simpson et al, 2016). Woody species composition was the slowest to respond to woody structural change, and we observed only minimal colonization of woody savanna species into the new grass layer of repeatedly burned thicket and forests over a decade of post-burn observations.

Together, these results suggest that extreme fires, when followed by subsequent fires, can trigger a biome switch from closed, fire-suppressing vegetation state to open, fire-dependent vegetation, structurally resembling savannas. This transition occurred on the time scale of a decade, which suggests that savanna-forest-thicket mosaics are likely dynamic even on the short-to-medium term (Rull *et al*, 2013, Beckett and Bond, 2019, but see Killeen et al, 2006). Repeated fire was crucial for maintaining an open canopy and allowing flammable grasses to colonise (see also Balch *et al.*, 2009, Silverio *et al.*, 2013), ultimately determining whether a biome shift from forest or thicket to savanna occurred.

In the transition from closed canopy vegetation states to savanna, initial grass colonization was dominated by *Panicum maximum,* which accumulated biomass rapidly. However, with 7 years of repeated fires, *Themeda triandra* gained in dominance in open-canopy systems. This replacement is consistent with the ecology of these two species: *Panicum maximum* (Paniceae) is often observed in relatively shaded savanna environments and is known to retain moisture longer into the dry season (Simpson et al, 2016), whereas *Themeda triandra* (Andropogoneae) prefers open, full-sun environments (Downing & Marshall, 1980, Kinyamario *et al.*, 1995) and is highly flammable (Simpson et al., 2016) but slower to colonise new environments (O’Connor & Everson, 1998).

On the time scale of a decade, however, novel savannas differed from old-growth savannas in their woody-layer composition (see also Veldman, 2016), and colonization of open habitats by savanna trees was slower than grass colonization. While forest canopies opened up readily, allowing grasses to colonise, the woody community remained depauperate of savanna trees (see also Veldman & Putz, 2011). Thickets differed from forests in that, even when burned repeatedly, they retained a tree community dominated by thicket species (probably resprouts). Meanwhile, forests that only burned once were colonised mostly by a unique pioneer assemblage. However, in novel savannas, frequent fires gradually depleted thicket and forest remnants, and we hypothesize that savanna woody species will eventually colonise (see also Charles-Dominique et al, 2015b). Further observation of previously closed-canopy areas that continue to experience frequent fires would allow this hypothesis to be tested.

Notably, we also found that forests could recover readily from a single extreme fire after a decade allowed the canopy to close and shade out grass fuels, thereby regaining its previous state. Forests likely recovered primarily from a seed bank, with an early-successional forest composed of rapidly growing pioneers (*i.e., Trema orientalis, Croton sylvaticus, Celtis africana)* and the clonally spreading liana *Dalbergia armata.* These species excluded light-demanding grasses, likely by shading and microclimate effects that also make the environment suitable for the establishment of climax forest species (see also Pammenter et al. 1985). Further work exploring a range of fire severities on the capability of a forest to recover would allow one to separate out the effects of an extreme fire with subsequent fires versus an extreme fire alone.

Thickets also recovered from a single fire readily but differed from forest in their post-fire recovery trajectory. While forests recovered via pioneer species germination, resulting in complete species turnover, thickets instead recovered via basal resprouting. Fire-driven tree mortality in thickets was low, as in savannas, contrasting with much higher tree mortality in forests (see also Ryan & Williams, 2011, Staver *et al.*, 2020). We predict that, on longer time scales, thickets and, by extension, other dry forests may be more resilient or tolerant of fires than wetter forests (see also Staal et al, 2020, Staver *et al,* 2020).

Our study documents a novel example of invasion of grass-fuelled fires into closed forest and thicket communities in an intact, ancient African savanna-forest-thicket mosaic. Our results offer a few key insights. Firstly, we show that extreme fires can allow savannas to invade forests even in the absence of invasive exotic grasses. Fires have been implicated in the expansion of the savanna biome from the late Miocene (Bond *et al.*, 2003, Keeley & Rundel, 2005, Scheiter *et al*, 2012, Behrensmeyer & Freeman, 2018), perhaps by gradually opening up forests and allowing shade-intolerant savanna grasses to spread at the forest edge (Schertzer & Staver,2018). Fires have also been implicated in constricting forest distributions in response to millennial scale changes in climate (Aleman *et al.*, 2019, Sato *et al*, 2021) – probably by increasing forest canopy openness, thereby facilitating grass invasion and eventually colonization by savanna trees. However, on-the-ground observations of this savannization process in action in intact landscapes are very few.

Secondly, we provide rare direct support for the longstanding hypothesis that runaway fire feedbacks, following an initial extreme fire, can result in the savannization of tropical forests (Cochrane, 1999) without additional anthropogenic drivers. In HiP, as in the Amazon, these extreme fires present a real problem. Their rarity makes them difficult to predict with any accuracy; however, we can be confident that extreme fire weather conditions are increasing in frequency as temperatures rise and aridity increases, making extreme fires an accelerating concern in savanna-forest mosaics, as they are elsewhere (Jolly *et al.*, 2015, Flannigan et al, 2013). Transitions from forest to savanna are indeed possible and likely to increase in potential extent in future.

The key difference between this study and work published under the savannization umbrella in the Neotropics revolves around the level of anthropogenic influence in these systems. Contemporary savannization in the Amazon is recorded in areas where exotic pasture grasses have invaded degraded forests, thereby increasing fuel loads and promoting subsequent fires (Silverio *et al*, 2013). However, paleoecological evidence suggests that savannization did occur in the Neotropics predating European colonization, probably as a results of fire in combination with low CO_2_ (Rull *et al*, 2013, Sato *et al*, 2021). It is interesting to note the potential impact the disappearance of Neotropical megafauna may have had on the probability of savannization occurring under non-anthropogenic conditions (Owen-Smith, 1987, Johnson, 2009). Recent work suggests that Neotropical grassy systems experienced a large increase in fire activity following the late-Quaternary collapse of large herbivore populations (Karp *et al.* 2021). Conversely, in African systems, the presence of herbivores (especially elephants) may open up the forest and thicket canopies along biome boundaries, allowing grass and therefore fire invasion and precipitating forest conversion to savanna. The role of megafauna in savannization processes has not been sufficiently investigated and offers an interesting avenue for future research.

Preventing or suppressing extreme fires is challenging in savannas, where fires are used as a management tool under less extreme conditions. In savannas, woody encroachment has long been a conservation concern, with widespread observations of increases in the woody component of savannas (Stevens *et al.*, 2017), particularly where fire is suppressed instead of being actively managed (Silva et al, 2008, Durigan, 2020). This trend is predicted to intensify as increasing CO_2_ favours C3 trees over C4 grasses (Higgins & Scheiter, 2012; Bond & Midgley, 2000, Moncrieff et al, 2014) – suggesting that, while forests are at increased risk of savannization, savannas are also at increased risk of forest expansion and woody encroachment. Widespread fire suppression is clearly not the solution for preventing extreme fires. Rather, we should aim instead for a rationalized fire management policy, allowing and encouraging fires in fire-dependent savannas but minimizing fires under extreme conditions, especially near fire-sensitive forests. Most crucially, we show that forests can recover readily from the effects of extreme fires, especially when an extreme event is not followed by subsequent fires, which may be easier to prevent.

## Acknowledgments

The financial assistance of SAEON towards this research is hereby acknowledged. Opinions expressed and conclusions arrived at are those of the author and are not necessarily to be attributed to SAEON.

## Author Contributions

HB, WJB, and ACS conceived of the idea. TCD and ACS contributed data. HB collected the data, designed, and performed the analyses with input from TCD and ACS. HB wrote the paper in consultation with WJB, ACS, and TCD.

