## Supplementary Tables and Figures for "Pathways of savannization in a mesic African savanna-forest mosaic following an extreme fire" (from another version of this preprint: version 2, 2021-09-21)

Supplementary Material

**Table S1.** Plot specific information for vegetation plots sampled in Hluhluwe-iMfolozi Park, displaying vegetation type, mean annual precipitation, years and the type of sampling that took place (T = Trees, G = Grasses), as well as the presence of closed canopy vegetation at various time points captured by aerial photography. Tree cover presence was calculated by Beckett and Bond (2019), where greater than 50% tree cover in a 10x10m cell was marked as tree cover presence.

| Plot ID | Source | Biome | Plot Type | Mean Annual Precipitation (mm)[1] | Sampling Done (G = Grass, T = Woody plants) 2007 | 2009 | 2012 | 2013 | 2014 | 2015 | Tree Cover Present (0 = Absent, 1 = Present)[2] 1937 | 1960 | 1975 | 1992 | 2006 | 2009 | 2013 |
|---|---|---|---|---|---|---|---|---|---|---|---|---|---|---|---|---|---|
| *t10* | *TCD* | *Thicket* | *Thicket* | 626 | | | | | | T | 0 | 0 | 1 | 1 | 1 | 1 | 1 |
| *t11* | *TCD* | *Thicket* | *Thicket* | 626 | | | | | | T | 0 | 0 | 1 | 1 | 1 | 1 | 1 |
| *t12* | *TCD* | *Thicket* | *Thicket* | 626 | | | | | | T | 0 | 0 | 1 | 1 | 1 | 1 | 1 |
| *t8* | *TCD* | *Thicket* | *Thicket* | 626 | | | | | | T | 0 | 0 | 1 | 1 | 1 | 1 | 1 |
| *t9* | *TCD* | *Thicket* | *Thicket* | 626 | | | | | | T | 0 | 0 | 1 | 1 | 1 | 1 | 1 |
| *t13* | *TCD* | *Thicket* | *Thicket* | 655 | | | | | | T | 0 | 0 | 1 | 1 | 1 | 1 | 1 |
| *t14* | *TCD* | *Thicket* | *Thicket* | 655 | | | | | | T | 0 | 0 | 1 | 1 | 1 | 1 | 1 |
| *t15* | *TCD* | *Thicket* | *Thicket* | 655 | | | | | | T | 0 | 0 | 1 | 1 | 1 | 1 | 1 |
| *t16* | *TCD* | *Thicket* | *Thicket* | 655 | | | | | | T | 0 | 0 | 1 | 1 | 1 | 1 | 1 |
| *t17* | *TCD* | *Thicket* | *Thicket* | 655 | | | | | | T | 0 | 0 | 1 | 1 | 1 | 1 | 1 |
| *t18* | *TCD* | *Thicket* | *Thicket* | 655 | | | | | | T | 0 | 0 | 1 | 1 | 1 | 1 | 1 |
| *t19* | *TCD* | *Thicket* | *Thicket* | 655 | | | | | | T | 0 | 0 | 1 | 1 | 1 | 1 | 1 |
| *t3* | *TCD* | *Thicket* | *Thicket* | 729 | | | | | | T | 0 | 0 | 1 | 1 | 1 | 1 | 1 |
| *t4* | *TCD* | *Thicket* | *Thicket* | 729 | | | | | | T | 0 | 0 | 1 | 1 | 1 | 1 | 1 |
| *t1* | *TCD* | *Thicket* | *Thicket* | 731 | | | | | | T | 0 | 0 | 1 | 1 | 1 | 1 | 1 |
| *t2* | *TCD* | *Thicket* | *Thicket* | 731 | | | | | | T | 0 | 0 | 1 | 1 | 1 | 1 | 1 |
| *t5* | *TCD* | *Thicket* | *Thicket* | 790 | | | | | | T | 0 | 1 | 1 | 1 | 1 | 1 | 1 |
| *f12* | *TCD* | *Forest* | *Forest* | 945 | | | | | | T | 1 | 1 | 1 | 1 | 1 | 1 | 1 |
| *f13* | *TCD* | *Forest* | *Forest* | 945 | | | | | | T | 1 | 1 | 1 | 1 | 1 | 1 | 1 |
| *f14* | *TCD* | *Forest* | *Forest* | 945 | | | | | | T | 1 | 1 | 1 | 1 | 1 | 1 | 1 |

*TCD – Charles-Dominique et al (2015), ACS&MC - Staver et al (2012) & Case & Staver (2017)*

*[1] Balfour and Howison, 2002, [2] Beckett and Bond, 2019*

**Table S1 cont.** Plot specific information for vegetation plots sampled in Hluhluwe-iMfolozi Park, displaying vegetation type, mean annual precipitation, years and the type of sampling that took place (T = Trees, G = Grasses), as well as the presence of closed canopy vegetation at various time points captured by aerial photography. Tree cover presence was calculated by Beckett and Bond (2019), where greater than 50% tree cover in a 10x10m cell was marked as tree cover presence.

| Plot | | | | Mean Annual Precipitation | Sampling Done (G = Grass, T = Woody plants) | | | | | | Tree Cover Present (0 = Absent, 1 = Present)[2] | | | | | | |
|---|---|---|---|---|---|---|---|---|---|---|---|---|---|---|---|---|---|
| ID | Source | Biome | Plot Type | (mm)[1] | 2007 | 2009 | 2012 | 2013 | 2014 | 2015 | 1937 | 1960 | 1975 | 1992 | 2006 | 2009 | 2013 |
| *f2* | *TCD* | *Forest* | *Forest* | 952 | | | | | | T | 1 | 1 | 1 | 1 | 1 | 1 | 1 |
| *f3* | *TCD* | *Forest* | *Forest* | 952 | | | | | | T | 1 | 1 | 1 | 1 | 1 | 1 | 1 |
| *f1* | *TCD* | *Forest* | *Forest* | 991 | | | | | | T | 1 | 1 | 1 | 1 | 1 | 1 | 1 |
| *f10* | *TCD* | *Forest* | *Forest* | 991 | | | | | | T | 1 | 1 | 1 | 1 | 1 | 1 | 1 |
| *f11* | *TCD* | *Forest* | *Forest* | 991 | | | | | | T | 1 | 1 | 1 | 1 | 1 | 1 | 1 |
| *f4* | *TCD* | *Forest* | *Forest* | 991 | | | | | | T | 1 | 1 | 1 | 1 | 1 | 1 | 1 |
| *f5* | *TCD* | *Forest* | *Forest* | 991 | | | | | | T | 1 | 1 | 1 | 1 | 1 | 1 | 1 |
| *f6* | *TCD* | *Forest* | *Forest* | 991 | | | | | | T | 1 | 1 | 1 | 1 | 1 | 1 | 1 |
| *f7* | *TCD* | *Forest* | *Forest* | 991 | | | | | | T | 1 | 1 | 1 | 1 | 1 | 1 | 1 |
| *f8* | *TCD* | *Forest* | *Forest* | 991 | | | | | | T | 1 | 1 | 1 | 1 | 1 | 1 | 1 |
| *f9* | *TCD* | *Forest* | *Forest* | 991 | | | | | | T | 1 | 1 | 1 | 1 | 1 | 1 | 1 |
| *s37* | *ACS&MC* | *Savanna* | *Savanna* | 658 | T | | | | T,G | | 0 | 0 | 0 | 0 | 0 | 0 | 0 |
| *s50* | *ACS&MC* | *Savanna* | *Savanna* | 663 | T | | | | | | 0 | 0 | 0 | 0 | 0 | 0 | 0 |
| *s29* | *ACS&MC* | *Savanna* | *Savanna* | 681 | T | | | | | | 0 | 0 | 0 | 0 | 0 | 0 | 0 |
| *s38* | *ACS&MC* | *Savanna* | *Savanna* | 686 | T | | | | T,G | | 0 | 0 | 0 | 0 | 0 | 0 | 0 |
| *s25* | *ACS&MC* | *Savanna* | *Savanna* | 694 | T | | | | | | 0 | 0 | 0 | 0 | 0 | 0 | 0 |
| *s44* | *ACS&MC* | *Savanna* | *Savanna* | 701 | T | | | | T,G | | 0 | 0 | 0 | 0 | 0 | 0 | 0 |
| *s49* | *ACS&MC* | *Savanna* | *Savanna* | 703 | T | | | | T,G | | 0 | 0 | 0 | 0 | 0 | 0 | 0 |
| *s26* | *ACS&MC* | *Savanna* | *Savanna* | 705 | T | | | | T,G | | 0 | 0 | 0 | 0 | 0 | 0 | 0 |
| *s23* | *ACS&MC* | *Savanna* | *Savanna* | 706 | T | | | | T,G | | 0 | 0 | 0 | 0 | 0 | 0 | 0 |

*TCD – Charles-Dominique et al (2015), ACS&MC - Staver et al (2012) & Case & Staver (2017)*

*[1] Balfour and Howison, 2002, [2] Beckett and Bond, 2019*

**Table S1 cont.** Plot specific information for vegetation plots sampled in Hluhluwe-iMfolozi Park, displaying vegetation type, mean annual precipitation, years and the type of sampling that took place (T = Trees, G = Grasses), as well as the presence of closed canopy vegetation at various time points captured by aerial photography. Tree cover presence was calculated by Beckett and Bond (2019), where greater than 50% tree cover in a 10x10m cell was marked as tree cover presence.

| Plot | | | Plot | Mean Annual Precipitation | Sampling Done (G = Grass, T = Woody plants) | | | | | | Tree Cover Present (0 = Absent, 1 = Present)[2] | | | | | | |
|---|---|---|---|---|---|---|---|---|---|---|---|---|---|---|---|---|---|
| ID | Source | Biome | Type | (mm)[1] | 2007 | 2009 | 2012 | 2013 | 2014 | 2015 | 1937 | 1960 | 1975 | 1992 | 2006 | 2009 | 2013 |
| *s24* | *ACS&MC* | *Savanna* | *Savanna* | 706 | T | | | | T,G | | 0 | 0 | 0 | 0 | 0 | 0 | 0 |
| *s27* | *ACS&MC* | *Savanna* | *Savanna* | 707 | T | | | | T,G | | 0 | 0 | 0 | 0 | 0 | 0 | 0 |
| *s39* | *ACS&MC* | *Savanna* | *Savanna* | 708 | T | | | | T,G | | 0 | 0 | 0 | 0 | 0 | 0 | 0 |
| *s40* | *ACS&MC* | *Savanna* | *Savanna* | 708 | T | | | | T,G | | 0 | 0 | 0 | 0 | 0 | 0 | 0 |
| *s18* | *ACS&MC* | *Savanna* | *Savanna* | 717 | T | | | | T,G | | 0 | 0 | 0 | 0 | 0 | 0 | 0 |
| *s28* | *ACS&MC* | *Savanna* | *Savanna* | 717 | T | | | | T,G | | 0 | 0 | 0 | 0 | 0 | 0 | 0 |
| *s19* | *ACS&MC* | *Savanna* | *Savanna* | 729 | T | | | | T,G | | 0 | 0 | 0 | 0 | 0 | 0 | 0 |
| *s20* | *ACS&MC* | *Savanna* | *Savanna* | 729 | T | | | | T,G | | 0 | 0 | 0 | 0 | 0 | 0 | 0 |
| *s41* | *ACS&MC* | *Savanna* | *Savanna* | 734 | T | | | | T,G | | 0 | 0 | 0 | 0 | 0 | 0 | 0 |
| *s45* | *ACS&MC* | *Savanna* | *Savanna* | 734 | T | | | | T,G | | 0 | 0 | 0 | 0 | 0 | 0 | 0 |
| *s46* | *ACS&MC* | *Savanna* | *Savanna* | 734 | T | | | | | | 0 | 0 | 0 | 0 | 0 | 0 | 0 |
| *s21* | *ACS&MC* | *Savanna* | *Savanna* | 738 | T | | | | T,G | | 0 | 0 | 0 | 0 | 0 | 0 | 0 |
| *s22* | *ACS&MC* | *Savanna* | *Savanna* | 738 | T | | | | T,G | | 0 | 0 | 0 | 0 | 0 | 0 | 0 |
| *s48* | *ACS&MC* | *Savanna* | *Savanna* | 747 | T | | | | | | 0 | 0 | 0 | 0 | 0 | 0 | 0 |
| *s13* | *ACS&MC* | *Savanna* | *Savanna* | 758 | T | | | | T,G | | 0 | 0 | 0 | 0 | 0 | 0 | 0 |
| *s10* | *ACS&MC* | *Savanna* | *Savanna* | 764 | T | | | | T,G | | 0 | 0 | 0 | 0 | 0 | 0 | 0 |
| *s11* | *ACS&MC* | *Savanna* | *Savanna* | 764 | T | | | | T,G | | 0 | 0 | 0 | 0 | 0 | 0 | 0 |
| *s12* | *ACS&MC* | *Savanna* | *Savanna* | 764 | T | | | | T,G | | 0 | 0 | 0 | 0 | 0 | 0 | 0 |
| *s17* | *ACS&MC* | *Savanna* | *Savanna* | 764 | T | | | | T,G | | 0 | 0 | 0 | 0 | 0 | 0 | 0 |
| *s8* | *ACS&MC* | *Savanna* | *Savanna* | 771 | T | | | | T,G | | 0 | 0 | 0 | 0 | 0 | 0 | 0 |

*TCD – Charles-Dominique et al (2015), ACS&MC - Staver et al (2012) & Case & Staver (2017)*

*[1] Balfour and Howison, 200,2[2] Beckett and Bond, 2019*

**Table S1 cont.** Plot specific information for vegetation plots sampled in Hluhluwe-iMfolozi Park, displaying vegetation type, mean annual precipitation, years and the type of sampling that took place (T = Trees, G = Grasses), as well as the presence of closed canopy vegetation at various time points captured by aerial photography. Tree cover presence was calculated by Beckett and Bond (2019), where greater than 50% tree cover in a 10x10m cell was marked as tree cover presence.

| Plot ID | Source | Biome | Plot Type | Mean Annual Precipitation (mm)[1] | Sampling Done (G = Grass, T = Woody plants) | | | | | | Tree Cover Present (0 = Absent, 1 = Present)[2] | | | | | | |
|---|---|---|---|---|---|---|---|---|---|---|---|---|---|---|---|---|---|
| | | | | | 2007 | 2009 | 2012 | 2013 | 2014 | 2015 | 1937 | 1960 | 1975 | 1992 | 2006 | 2009 | 2013 |
| *s9* | *ACS&MC* | *Savanna* | *Savanna* | 771 | T | | | | T,G | | 0 | 0 | 0 | 0 | 0 | 0 | 0 |
| *s30* | *ACS&MC* | *Savanna* | *Savanna* | 771 | T | | | | T,G | | 0 | 0 | 0 | 0 | 0 | 0 | 0 |
| *s51* | *ACS&MC* | *Savanna* | *Savanna* | 771 | T | | | | T,G | | 0 | 0 | 0 | 0 | 0 | 0 | 0 |
| *s15* | *ACS&MC* | *Savanna* | *Savanna* | 782 | T | | | | T,G | | 0 | 0 | 0 | 0 | 0 | 0 | 0 |
| *s16* | *ACS&MC* | *Savanna* | *Savanna* | 782 | T | | | | T,G | | 0 | 0 | 0 | 0 | 0 | 0 | 0 |
| *s14* | *ACS&MC* | *Savanna* | *Savanna* | 785 | T | | | | T,G | | 0 | 0 | 0 | 0 | 0 | 0 | 0 |
| *s6* | *ACS&MC* | *Savanna* | *Savanna* | 789 | T | | | | T,G | | 0 | 0 | 0 | 0 | 0 | 0 | 0 |
| *s7* | *ACS&MC* | *Savanna* | *Savanna* | 789 | T | | | | T,G | | 0 | 0 | 0 | 0 | 0 | 0 | 0 |
| *s1* | *ACS&MC* | *Savanna* | *Savanna* | 790 | T | | | | T,G | | 0 | 0 | 0 | 0 | 0 | 0 | 0 |
| *s2* | *ACS&MC* | *Savanna* | *Savanna* | 790 | T | | | | T,G | | 0 | 0 | 0 | 0 | 0 | 0 | 0 |
| *s42* | *ACS&MC* | *Savanna* | *Savanna* | 798 | T | | | | | | 0 | 0 | 0 | 0 | 0 | 0 | 0 |
| *s43* | *ACS&MC* | *Savanna* | *Savanna* | 798 | T | | | | T,G | | 0 | 0 | 0 | 0 | 0 | 0 | 0 |
| *s47* | *ACS&MC* | *Savanna* | *Savanna* | 803 | T | | | | | | 0 | 0 | 0 | 0 | 0 | 0 | 0 |
| *s32* | *ACS&MC* | *Savanna* | *Savanna* | 807 | T | | | | T,G | | 0 | 0 | 0 | 0 | 0 | 0 | 0 |
| *s33* | *ACS&MC* | *Savanna* | *Savanna* | 807 | T | | | | T,G | | 0 | 0 | 0 | 0 | 0 | 0 | 0 |
| *s31* | *ACS&MC* | *Savanna* | *Savanna* | 812 | T | | | | T,G | | 0 | 0 | 0 | 0 | 0 | 0 | 0 |
| *s4* | *ACS&MC* | *Savanna* | *Savanna* | 820 | T | | | | T,G | | 0 | 0 | 0 | 0 | 0 | 0 | 0 |
| *s5* | *ACS&MC* | *Savanna* | *Savanna* | 820 | T | | | | T,G | | 0 | 0 | 0 | 0 | 0 | 0 | 0 |
| *s3* | *ACS&MC* | *Savanna* | *Savanna* | 828 | T | | | | T,G | | 0 | 0 | 0 | 0 | 0 | 0 | 0 |
| *s34* | *ACS&MC* | *Savanna* | *Savanna* | 840 | T | | | | T,G | | 0 | 0 | 0 | 0 | 0 | 0 | 0 |

*TCD – Charles-Dominique et al (2015), ACS&MC - Staver et al (2012) & Case & Staver (2017)*

*[1] Balfour and Howison, 2002, [2] Beckett and Bond, 2019*

**Table S1 cont.** Plot specific information for vegetation plots sampled in Hluhluwe-iMfolozi Park, displaying vegetation type, mean annual precipitation, years and the type of sampling that took place (T = Trees, G = Grasses), as well as the presence of closed canopy vegetation at various time points captured by aerial photography. Tree cover presence was calculated by Beckett and Bond (2019), where greater than 50% tree cover in a 10x10m cell was marked as tree cover presence.

| Plot ID | Source | Biome | Plot Type | Mean Annual Precipitation (mm)[1] | Sampling Done (G = Grass, T = Woody plants) | | | | | | Tree Cover Present (0 = Absent, 1 = Present)[2] | | | | | | |
|---|---|---|---|---|---|---|---|---|---|---|---|---|---|---|---|---|---|
| | | | | | 2007 | 2009 | 2012 | 2013 | 2014 | 2015 | 1937 | 1960 | 1975 | 1992 | 2006 | 2009 | 2013 |
| *s35* | *ACS&MC* | *Savanna* | *Savanna* | 840 | T | | | | T,G | | 0 | 0 | 0 | 0 | 0 | 0 | 0 |
| *s36* | *ACS&MC* | *Savanna* | *Savanna* | 866 | T | | | | T,G | | 0 | 0 | 0 | 0 | 0 | 0 | 0 |
| *f19* | *This study* | *Forest* | *Forest* | 945 | | | | | T,G | | 1 | 1 | 1 | 1 | 1 | 1 | 1 |
| *f15* | *This study* | *Forest* | *Forest* | 952 | | | | | T,G | | 1 | 1 | 1 | 1 | 1 | 1 | 1 |
| *f16* | *This study* | *Forest* | *Forest* | 952 | | | | | T,G | | 1 | 1 | 1 | 1 | 1 | 1 | 1 |
| *f17* | *This study* | *Forest* | *Forest* | 991 | | | | | T,G | | 1 | 1 | 1 | 1 | 1 | 1 | 1 |
| *f18* | *This study* | *Forest* | *Forest* | 991 | | | | | T,G | | 1 | 1 | 1 | 1 | 1 | 1 | 1 |
| *rf22* | *This study* | *Forest* | *Frequently-burnt Forest* | 729 | | G | G | | | T,G | 1 | 1 | 1 | 1 | 1 | 0 | 0 |
| *rf15* | *This study* | *Forest* | *Frequently-burnt Forest* | 850 | | G | G | | | T,G | 1 | 1 | 1 | 1 | 1 | 0 | 0 |
| *rf16* | *This study* | *Forest* | *Frequently-burnt Forest* | 850 | | G | G | | | T,G | 1 | 1 | 1 | 1 | 1 | 0 | 0 |
| *rf17* | *This study* | *Forest* | *Frequently-burnt Forest* | 850 | | G | G | | | T,G | 1 | 1 | 1 | 1 | 1 | 0 | 0 |
| *rf18* | *This study* | *Forest* | *Frequently-burnt Forest* | 850 | | G | G | | | T,G | 1 | 1 | 1 | 1 | 1 | 0 | 0 |
| *rf19* | *This study* | *Forest* | *Frequently-burnt Forest* | 866 | | G | G | | | T,G | 1 | 1 | 1 | 1 | 1 | 0 | 0 |
| *rf4* | *This study* | *Forest* | *Frequently-burnt Forest* | 913 | | | | | | T,G | 1 | 1 | 1 | 1 | 1 | 0 | 0 |
| *rf6* | *This study* | *Forest* | *Frequently-burnt Forest* | 913 | | | | | | T,G | 1 | 1 | 1 | 1 | 1 | 0 | 0 |
| *rf7* | *This study* | *Forest* | *Frequently-burnt Forest* | 913 | | | | | | T,G | 1 | 1 | 1 | 1 | 1 | 0 | 0 |
| *rf9* | *This study* | *Forest* | *Frequently-burnt Forest* | 913 | | | | | | T,G | 1 | 1 | 1 | 1 | 1 | 0 | 0 |
| *rf3* | *This study* | *Forest* | *Frequently-burnt Forest* | 915 | | | | | | T,G | 1 | 1 | 1 | 1 | 1 | 0 | 0 |
| *rf20* | *This study* | *Forest* | *Frequently-burnt Forest* | 917 | | G | G | | | T,G | 1 | 1 | 1 | 1 | 1 | 0 | 0 |
| *rf21* | *This study* | *Forest* | *Frequently-burnt Forest* | 917 | | G | G | | | T,G | 1 | 1 | 1 | 1 | 1 | 0 | 0 |

*TCD – Charles-Dominique et al (2015), ACS&MC - Staver et al (2012) & Case & Staver (2017)*

*[1] Balfour and Howison, 2002, [2] Beckett and Bond, 2019*

**Table S1 cont.** Plot specific information for vegetation plots sampled in Hluhluwe-iMfolozi Park, displaying vegetation type, mean annual precipitation, years and the type of sampling that took place (T = Trees, G = Grasses), as well as the presence of closed canopy vegetation at various time points captured by aerial photography. Tree cover presence was calculated by Beckett and Bond (2019), where greater than 50% tree cover in a 10x10m cell was marked as tree cover presence.

| Plot ID | Source | Biome | Plot Type | Mean Annual Precipitation (mm)[1] | Sampling Done (G = Grass, T = Woody plants) | | | | | | Tree Cover Present (0 = Absent, 1 = Present)[2] | | | | | | |
|---|---|---|---|---|---|---|---|---|---|---|---|---|---|---|---|---|---|
| | | | | | 2007 | 2009 | 2012 | 2013 | 2014 | 2015 | 1937 | 1960 | 1975 | 1992 | 2006 | 2009 | 2013 |
| *rf5* | *This study* | *Forest* | *Frequently-burnt Forest* | 926 | | | | | | T,G | 1 | 1 | 1 | 1 | 1 | 0 | 0 |
| *rf13* | *This study* | *Forest* | *Recovering Forest* | 900 | | | | | T,G | T,G | 1 | 1 | 1 | 1 | 1 | 0 | 0 |
| *rf14* | *This study* | *Forest* | *Recovering Forest* | 900 | | | | | T,G | T,G | 1 | 1 | 1 | 1 | 1 | 0 | 0 |
| *rf8* | *This study* | *Forest* | *Recovering Forest* | 913 | | | | | T,G | | 1 | 1 | 1 | 1 | 1 | 0 | 0 |
| *rf1* | *This study* | *Forest* | *Recovering Forest* | 991 | | | | | | T,G | 1 | 1 | 1 | 1 | 0 | 0 | 0 |
| *rf10* | *This study* | *Forest* | *Recovering Forest* | 991 | | | | T,G | | | 1 | 1 | 1 | 1 | 0 | 0 | 0 |
| *rf11* | *This study* | *Forest* | *Recovering Forest* | 991 | | | | T,G | | | 1 | 1 | 1 | 1 | 0 | 0 | 0 |
| *rf12* | *This study* | *Forest* | *Recovering Forest* | 991 | | | | T,G | | | 1 | 1 | 1 | 1 | 0 | 0 | 0 |
| *rf2* | *This study* | *Forest* | *Recovering Forest* | 991 | | | | | | T,G | 1 | 1 | 1 | 1 | 0 | 0 | 0 |
| *rt6* | *This study* | *Thicket* | *Frequently-burnt Thicket* | 694 | | G | G | | | T,G | 0 | 1 | 1 | 1 | 1 | 0 | 0 |
| *rt7* | *This study* | *Thicket* | *Frequently-burnt Thicket* | 694 | | G | G | | | T,G | 0 | 1 | 1 | 1 | 1 | 0 | 0 |
| *rt10* | *This study* | *Thicket* | *Frequently-burnt Thicket* | 706 | | G | G | | | T,G | 0 | 0 | 1 | 1 | 1 | 0 | 0 |
| *rt11* | *This study* | *Thicket* | *Frequently-burnt Thicket* | 706 | | G | G | | | T,G | 0 | 0 | 1 | 1 | 1 | 0 | 0 |
| *rt8* | *This study* | *Thicket* | *Frequently-burnt Thicket* | 706 | | G | G | | | T,G | 0 | 0 | 1 | 1 | 1 | 0 | 0 |
| *rt9* | *This study* | *Thicket* | *Frequently-burnt Thicket* | 706 | | G | G | | | T,G | 0 | 0 | 1 | 1 | 1 | 0 | 0 |
| *rt1* | *This study* | *Thicket* | *Frequently-burnt Thicket* | 729 | | G | G | | | T,G | 0 | 0 | 1 | 1 | 1 | 0 | 0 |
| *rt12* | *This study* | *Thicket* | *Frequently-burnt Thicket* | 731 | | G | G | | | T,G | 0 | 0 | 0 | 1 | 1 | 0 | 0 |
| *rt2* | *This study* | *Thicket* | *Frequently-burnt Thicket* | 742 | | G | G | | | T,G | 0 | 0 | 0 | 1 | 1 | 0 | 0 |
| *rt3* | *This study* | *Thicket* | *Frequently-burnt Thicket* | 742 | | G | G | | | T,G | 0 | 0 | 0 | 1 | 1 | 0 | 0 |
| *rt4* | *This study* | *Thicket* | *Frequently-burnt Thicket* | 812 | | G | G | | | T,G | 0 | 0 | 0 | 1 | 1 | 0 | 0 |

*TCD – Charles-Dominique et al (2015), ACS&MC - Staver et al (2012) & Case & Staver (2017)*

*[1] Balfour and Howison, 2002, [2] Beckett and Bond, 2019*

**Table S1 cont.** Plot specific information for vegetation plots sampled in Hluhluwe-iMfolozi Park, displaying vegetation type, mean annual precipitation, years and the type of sampling that took place (T = Trees, G = Grasses), as well as the presence of closed canopy vegetation at various time points captured by aerial photography. Tree cover presence was calculated by Beckett and Bond (2019), where greater than 50% tree cover in a 10x10m cell was marked as tree cover presence.

| Plot ID | Source | Biome | Plot Type | Mean Annual Precipitation (mm)[1] | Sampling Done (G = Grass, T = Woody plants) | | | | | | Tree Cover Present (0 = Absent, 1 = Present)[2] | | | | | | |
|---|---|---|---|---|---|---|---|---|---|---|---|---|---|---|---|---|---|
| | | | | | 2007 | 2009 | 2012 | 2013 | 2014 | 2015 | 1937 | 1960 | 1975 | 1992 | 2006 | 2009 | 2013 |
| *rt5* | *This study* | *Thicket* | *Frequently-burnt Thicket* | 812 | | G | G | | | T,G | 0 | 0 | 1 | 1 | 1 | 0 | 0 |
| *t38* | *This study* | *Thicket* | *Thicket* | 729 | G | | G | | T,G | | 0 | 0 | 1 | 1 | 1 | 1 | 1 |
| *t45* | *This study* | *Thicket* | *Thicket* | 729 | | | | | T,G | | 0 | 0 | 0 | 1 | 1 | 1 | 1 |
| *t40* | *This study* | *Thicket* | *Thicket* | 731 | G | | G | | T,G | | 0 | 0 | 1 | 1 | 1 | 1 | 1 |
| *t41* | *This study* | *Thicket* | *Thicket* | 731 | G | | G | | | | 0 | 0 | 1 | 1 | 1 | 1 | 0 |
| *t39* | *This study* | *Thicket* | *Thicket* | 746 | G | | G | | | | 0 | 0 | 1 | 1 | 1 | 1 | 0 |
| *t44* | *This study* | *Thicket* | *Thicket* | 771 | | | | | T,G | | 0 | 1 | 1 | 1 | 1 | 1 | 1 |
| *t43* | *This study* | *Thicket* | *Thicket* | 790 | | | | | T,G | | 0 | 1 | 1 | 1 | 1 | 1 | 1 |
| *t42* | *This study* | *Thicket* | *Thicket* | 828 | | | | | T,G | | 0 | 1 | 1 | 1 | 1 | 1 | 1 |

*TCD – Charles-Dominique et al (2015), ACS&MC – Staver et al (2012) & Case & Staver (2017)*
*[1] Balfour and Howison, 2002, [2] Beckett and Bond, 2019*

**Table S2.** Median Stem Density for vegetation types based on the plots sampled in Hluhluwe-iMfolozi Park, South Africa. Values shown represent the total density for all size classes, and individual size class medians. Values in parentheses are the interquartile ranges.

| Biome | Vegetation Type | N | Median Stem Density | | | | |
|---|---|---|---|---|---|---|---|
| | | | Total | 0.5 - 2m | 2 - 5m | 5 - 10m | > 10m |
| Forest | *Intact* | 19 | 107 (87, 130) | 30 (15, 32) | 27 (16, 37) | 29 (22, 42) | 27 (20, 38) |
| | *Frequently Burnt* | 14 | 150 (127, 164) | 138 (120, 149) | 10 (2, 21) | 0 (0, 1) | 0 |
| | *Recovering (6 years)* | 3 | 167 (166, 172) | 118 (110, 120) | 38 (36, 48) | 16 (10, 16) | 0 |
| | *Recovering (9 years)* | 3 | 170 (132, 172) | 75 (64, 84) | 58 (46, 69) | 15 (10, 20) | 0 |
| | *Recovering (11 years)* | 2 | 420 (398, 443) | 168 (149, 188) | 212 (211, 212) | 37 (35, 39) | 4 (2, 5) |
| Savanna | *2007* | 51 | 178 (124, 234) | 166 (117, 228) | 6 (2, 10) | 0 (0, 1) | 0 |
| | *2014* | 44 | 186 (118, 230) | 170 (108, 223) | 3 (1, 9) | 0 | 0 |
| Thicket | *Intact* | 23 | 278 (202, 336) | 116 (98, 140) | 93 (78, 108) | 30 (13, 60) | 2 (0, 14) |
| | *Frequently Burnt* | 12 | 137 (89, 156) | 130 (88, 154) | 2 (0, 6) | 0 | 0 |

**Table S3**. Vegetation type comparisons of cumulative length of woody vegetation within different size classes, showing the results of the MANOVA, and the *f* and *p*-values from univariate ANOVAs on the size classes within those comparisons.

| Vegetation Types | MANOVA | Univariate ANOVA | | | |
|---|---|---|---|---|---|
| | | Size Class | Df | F Value | Pr(>F) |
| *Savanna 2007 – Savanna 2014* | *Pillai's Trace = 0.039, Approx* $F_{(3,91)} = 1.238$, $p = 0.301$ | *0.5 - 2m* | 1 | 0.3739 | 0.542 |
| | | *2 - 5m* | 1 | 0.2304 | 0.632 |
| | | *5 - 10m* | 1 | 2.6188 | 0.109 |
| | | *> 10m* | 1 | NA | NA |
| *Intact Forest - Savanna* | *Pillai's Trace = 0.88, Approx* $F_{(4,58)} = 109.69$, $p < 0.001$ | *0.5 - 2m* | 1 | 72.16 | < 0.0001 |
| | | *2 - 5m* | 1 | 12.703 | < 0.0001 |
| | | *5 - 10m* | 1 | 189.58 | < 0.0001 |
| | | *> 10m* | 1 | 247.33 | < 0.0001 |
| *Intact Thicket - Savanna* | *Pillai's Trace = 0.75, Approx* $F_{(4,62)} = 47.484$, $p < 0.001$ | *0.5 - 2m* | 1 | 8.3367 | 0.005 |
| | | *2 - 5m* | 1 | 148.02 | < 0.0001 |
| | | *5 - 10m* | 1 | 80.073 | < 0.0001 |
| | | *> 10m* | 1 | 29.384 | < 0.0001 |
| *Intact Forest - Intact Thicket* | *Pillai's Trace = 0.83, Approx* $F_{(4,37)} = 45.622$, $p < 0.001$ | *0.5 - 2m* | 1 | 126.63 | < 0.0001 |
| | | *2 - 5m* | 1 | 56.038 | < 0.0001 |
| | | *5 - 10m* | 1 | 0.7285 | 0.398 |
| | | *> 10m* | 1 | 44.77 | < 0.0001 |
| *Intact Forest - Frequently Burnt Forest* | *Pillai's Trace = 0.89, Approx* $F_{(4,28)} = 58.734$, $p < 0.001$ | *0.5 - 2m* | 1 | 141.57 | < 0.0001 |
| | | *2 - 5m* | 1 | 2.7457 | 0.108 |
| | | *5 - 10m* | 1 | 54.543 | < 0.0001 |
| | | *> 10m* | 1 | 75.938 | < 0.0001 |

**Table S3 cont**. Vegetation type comparisons of cumulative length of woody vegetation within different size classes, showing the results of the MANOVA, and the *f* and *p*-values from univariate ANOVAs on the size classes within those comparisons.

| Vegetation Types | MANOVA | Univariate ANOVA | | | |
|---|---|---|---|---|---|
| | | Size Class | Df | F Value | Pr(>F) |
| *Frequently Burnt Forest - Savanna* | *Pillai's Trace = 0.20, Approx $F_{(4,53)}$ = 3.3665, p = 0.0158* | *0.5 - 2m* | 1 | 3.8498 | 0.055 |
| | | *2 - 5m* | 1 | 1.5379 | 0.220 |
| | | *5 - 10m* | 1 | 5.6093 | 0.021 |
| | | *> 10m* | 1 | 3.2679 | 0.076 |
| *Intact Thicket - Frequently Burnt Thicket* | *Pillai's Trace = 0.76, Approx $F_{(4,30)}$ = 23.406, p < 0.001* | *0.5 - 2m* | 1 | 0.4196 | 0.522 |
| | | *2 - 5m* | 1 | 67.157 | < 0.0001 |
| | | *5 - 10m* | 1 | 21.281 | < 0.0001 |
| | | *> 10m* | 1 | 7.5985 | 0.009 |
| *Frequently Burnt Thicket - Savanna* | *Pillai's Trace = 0.15, Approx $F_{(4,51)}$ = 2.2415, p = 0.0774* | *0.5 - 2m* | 1 | 2.4965 | 0.120 |
| | | *2 - 5m* | 1 | 0.5708 | 0.453 |
| | | *5 - 10m* | 1 | 0.0449 | 0.833 |
| | | *> 10m* | 1 | 3.8571 | 0.055 |
| *Frequently Burnt Forest - Frequently Burnt Thicket* | *Pillai's Trace = 0.16, Approx $F_{(4,21)}$ = 0.95975, p = 0.45* | *0.5 - 2m* | 1 | 0.0355 | 0.852 |
| | | *2 - 5m* | 1 | 3.5479 | 0.072 |
| | | *5 - 10m* | 1 | 1.7456 | 0.199 |
| | | *> 10m* | 1 | 0.0119 | 0.914 |
| *Frequently Burnt Forest - Once Burnt Forest 6yrs* | *Pillai's Trace = 0.61, Approx $F_{(4,12)}$ = 4.7211, p = 0.016* | *0.5 - 2m* | 1 | 0.4221 | 0.526 |
| | | *2 - 5m* | 1 | 3.886 | 0.0674 |
| | | *5 - 10m* | 1 | 21.956 | < 0.001 |
| | | *> 10m* | 1 | 0.2036 | 0.6583 |

**Table S3 cont**. Vegetation type comparisons of cumulative length of woody vegetation within different size classes, showing the results of the MANOVA, and the *f* and *p*-values from univariate ANOVAs on the size classes within those comparisons.

| Vegetation Types | MANOVA | Univariate ANOVA | | | |
|---|---|---|---|---|---|
| | | Size Class | Df | F Value | Pr(>F) |
| *Intact Forest – Once burnt Forest 6yrs* | *Pillai's Trace = 0.902, Approx* $F_{(4,17)}$ *= 38.902, p = < 0.0001* | *0.5 - 2m* | 1 | 158.48 | < 0.0001 |
| | | *2 - 5m* | 1 | 4.1919 | 0.054 |
| | | *5 - 10m* | 1 | 4.8527 | 0.0395 |
| | | *> 10m* | 1 | 15.833 | < 0.001 |
| *Savanna - Once Burnt Forest 6yrs* | *Pillai's Trace = 0.765, Approx* $F_{(3,43)}$ *= 46.72, p < 0.0001* | *0.5 - 2m* | 1 | 1.5881 | 0.2141 |
| | | *2 - 5m* | 1 | 8.3046 | 0.00604 |
| | | *5 - 10m* | 1 | 126.23 | < 0.0001 |
| | | *> 10m* | 1 | NA | NA |
| *Frequently Burnt Forest - Once Burnt Forest 9yrs* | *Pillai's Trace = 0.716, Approx* $F_{(4,12)}$ *= 7.5689, p = 0.0028* | *0.5 - 2m* | 1 | 6.5341 | 0.022 |
| | | *2 - 5m* | 1 | 8.423 | 0.0110 |
| | | *5 - 10m* | 1 | 26.297 | 0.00012 |
| | | *> 10m* | 1 | 0.204 | 0.658 |
| *Intact Forest – Once burnt Forest 9yrs* | *Pillai's Trace = 0.761, Approx* $F_{(4,17)}$ *= 13.499, p < 0.0001* | *0.5 - 2m* | 1 | 37.751 | < 0.0001 |
| | | *2 - 5m* | 1 | 12.352 | 0.00218 |
| | | *5 - 10m* | 1 | 3.641 | 0.0708 |
| | | *> 10m* | 1 | 15.833 | 0.0007 |
| *Savanna - Once Burnt Forest 9yrs* | *Pillai's Trace = 0.767, Approx* $F_{(3,43)}$ *= 47.296, p < 0.0001* | *0.5 - 2m* | 1 | 4.977 | 0.2141 |
| | | *2 - 5m* | 1 | 16.096 | 0.00022 |
| | | *5 - 10m* | 1 | 133.29 | < 0.0001 |
| | | *> 10m* | 1 | NA | NA |

**Table S4**. PERMANOVA and Pairwise PERMANOVAs on a Bray-Curtis dissimilarity matrix of the abundance of species types (Forest, Savanna, Thicket & Pioneer) combined across all size classes for the different vegetation types in Hluhluwe-iMfolozi Park. Adjusted P values, calculated using the procedure of Benjamini and Hochberg (1995) are presented for the pairwise comparisons.

| **Source** | **d.f.** | **SS** | **R2** | **F** | **P** |
|---|---|---|---|---|---|
| *Vegetation Type* | 8 | 23.0127 | 0.814 | 89.643 | **0.001** |
| *Residual* | 164 | 5.2626 | 0.186 | | |
| *Total* | 172 | 28.2754 | | | |

| **Pair-wise Comparisons** | | **F** | **R2** | **P** | **adj. P** |
|---|---|---|---|---|---|
| *Savanna 2007* | *Savanna 2014* | 1.270 | 0.013 | 0.294 | 0.3000 |
| *Savanna 2007* | *Intact Thicket* | 136.009 | 0.648 | **0.001** | **0.0016** |
| *Savanna 2007* | *Frequently Burnt Thicket* | 50.752 | 0.454 | **0.001** | **0.0016** |
| *Savanna 2007* | *Intact Forest* | 232.410 | 0.774 | **0.001** | **0.0016** |
| *Savanna 2007* | *Frequently Burnt Forest* | 63.736 | 0.503 | **0.001** | **0.0016** |
| *Savanna 2007* | *Once Burnt Forest (6 years)* | 32.438 | 0.384 | **0.001** | **0.0016** |
| *Savanna 2007* | *Once Burnt Forest (9 years)* | 29.947 | 0.365 | **0.001** | **0.0016** |
| *Intact Forest* | *Intact Thicket* | 152.319 | 0.784 | **0.001** | **0.0016** |
| *Intact Forest* | *Frequently Burnt Forest* | 61.399 | 0.665 | **0.001** | **0.0016** |
| *Intact Forest* | *Once Burnt Forest (6 years)* | 33.061 | 0.623 | **0.001** | **0.0016** |
| *Intact Forest* | *Once Burnt Forest (9 years)* | 16.864 | 0.457 | **0.001** | **0.0016** |
| *Frequently Burnt Forest* | *Frequently Burnt Thicket* | 6.890 | 0.223 | **0.001** | **0.0016** |
| *Frequently Burnt Forest* | *Once Burnt Forest (6 years)* | 1.984 | 0.117 | 0.102 | 0.1080 |
| *Frequently Burnt Forest* | *Once Burnt Forest (9 years)* | 3.278 | 0.179 | **0.024** | **0.0279** |
| *Once Burnt Forest (6 years)* | *Once Burnt Forest (9 years)* | 0.937 | 0.190 | 0.300 | 0.3000 |
| *Intact Thicket* | *Frequently Burnt Thicket* | 28.863 | 0.452 | **0.001** | **0.0016** |

**Table S5**. PERMANOVA and Pairwise PERMANOVAs on a Bray-Curtis dissimilarity matrix of the abundance of species types (Forest, Savanna, Thicket & Pioneer) within each size class for the different vegetation types in Hluhluwe-iMfolozi Park. Adjusted P values, calculated using the procedure of Benjamini and Hochberg (1995) are presented for the pairwise comparisons.

| Source | d.f. | SS | R2 | F | P |
|---|---|---|---|---|---|
| *Vegetation Type* | 8 | 27.744 | 0.761 | 65.118 | **0.001** |
| *Residual* | 164 | 8.734 | 0.239 | | |
| *Total* | 172 | 36.478 | | | |

| Pair-wise Comparisons | | F | R2 | P | adj. P |
|---|---|---|---|---|---|
| *Savanna 2007* | *Savanna 2014* | 0.996 | 0.011 | 0.351 | 0.3510 |
| *Savanna 2007* | *Intact Thicket* | 119.675 | 0.618 | **0.001** | **0.0016** |
| *Savanna 2007* | *Frequently Burnt Thicket* | 40.875 | 0.401 | **0.001** | **0.0016** |
| *Savanna 2007* | *Intact Forest* | 169.307 | 0.713 | **0.001** | **0.0016** |
| *Savanna 2007* | *Frequently Burnt Forest* | 48.943 | 0.437 | **0.001** | **0.0016** |
| *Savanna 2007* | *Once Burnt Forest (6 years)* | 25.034 | 0.325 | **0.001** | **0.0016** |
| *Savanna 2007* | *Once Burnt Forest (9 years)* | 24.522 | 0.320 | **0.001** | **0.0016** |
| *Intact Forest* | *Intact Thicket* | 117.207 | 0.736 | **0.001** | **0.0016** |
| *Intact Forest* | *Frequently Burnt Forest* | 53.410 | 0.633 | **0.001** | **0.0016** |
| *Intact Forest* | *Once Burnt Forest (6 years)* | 28.749 | 0.590 | **0.001** | **0.0016** |
| *Intact Forest* | *Once Burnt Forest (9 years)* | 22.914 | 0.534 | **0.001** | **0.0016** |
| *Frequently Burnt Forest* | *Frequently Burnt Thicket* | 6.595 | 0.216 | **0.001** | **0.0016** |
| *Frequently Burnt Forest* | *Once Burnt Forest (6 years)* | 2.403 | 0.138 | **0.036** | **0.0405** |
| *Frequently Burnt Forest* | *Once Burnt Forest (9 years)* | 5.136 | 0.255 | **0.001** | **0.0016** |
| *Once Burnt Forest (6 years)* | *Once Burnt Forest (9 years)* | 1.446 | 0.266 | 0.300 | 0.3086 |
| *Intact Thicket* | *Frequently Burnt Thicket* | 32.845 | 0.484 | **0.001** | **0.0016** |

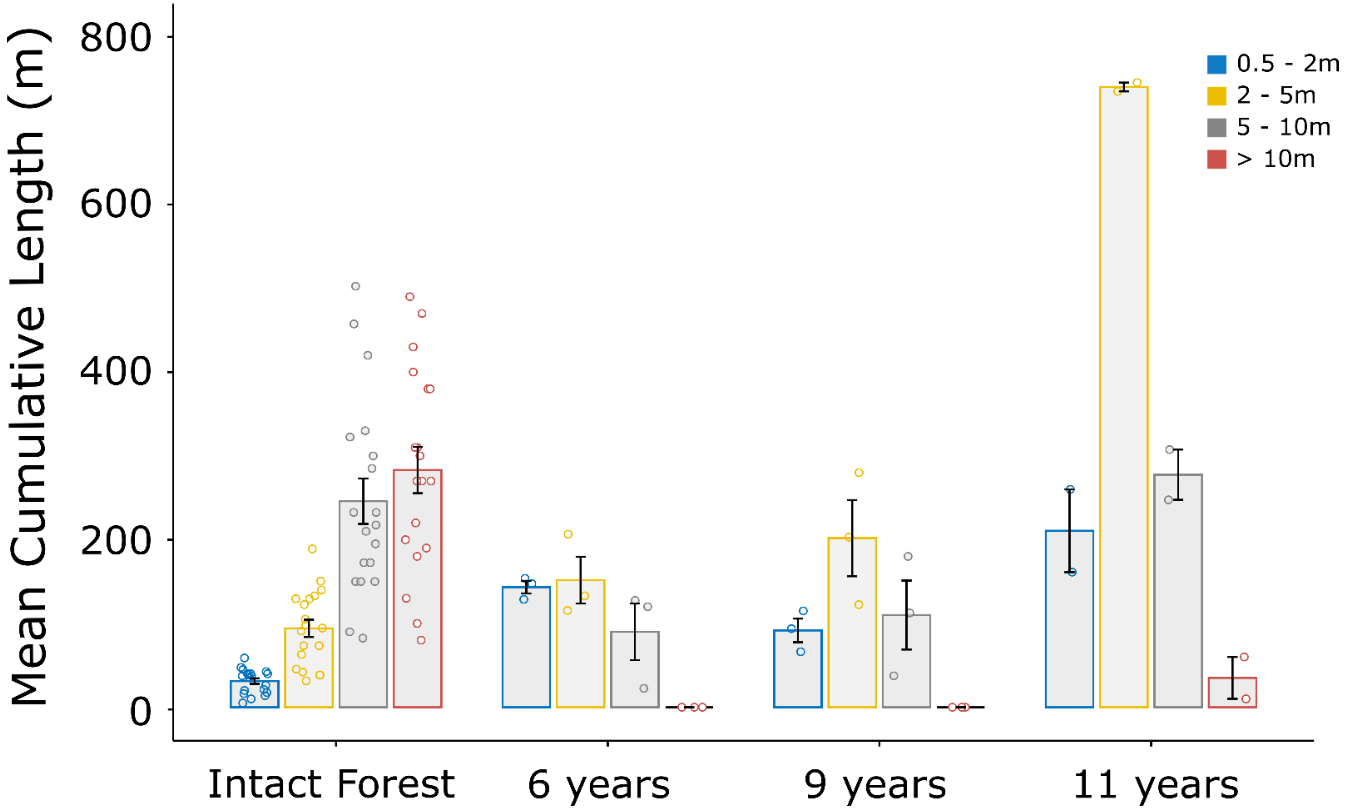


**Appendix Figure S1.** Mean cumulative length of all tree species within the four size classes in intact forest and after the extreme fire event in once-burnt forest at different time periods. Points represent individual transects. The 9- and 11-years plots include three and two sites respectively as one plot was burnt 10 years after the 2004 fire and was removed. These three plots covered more than 80% of the forest area burnt in 2004, hence the low sample size.

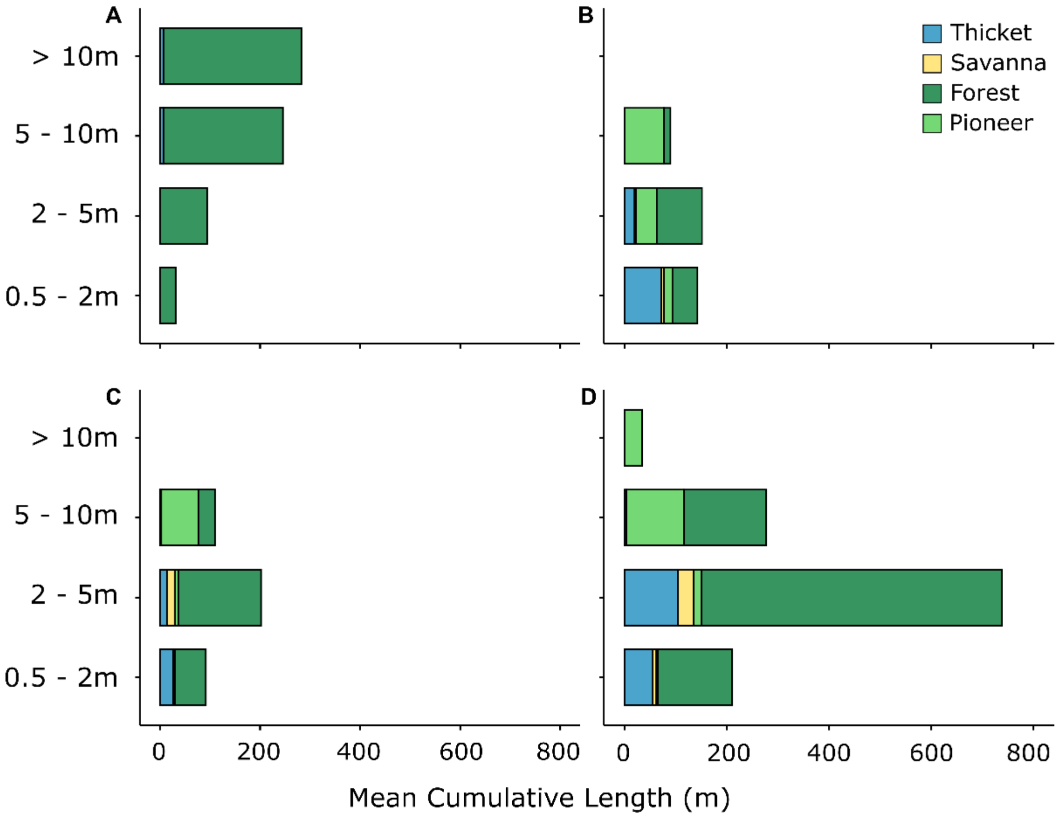


**Appendix Figure S2.** Initial and post fire composition and structure of intact forest (A) and once burnt forest after six (B), nine (C) and 11 years (D). Panels display the average cumulative length of tree species within each size class for the different time periods before and after the extreme fire event. Colours indicate the biome in which these species are predominantly found. Pioneer species are those which were not found in transects sampled before the extreme fire event.

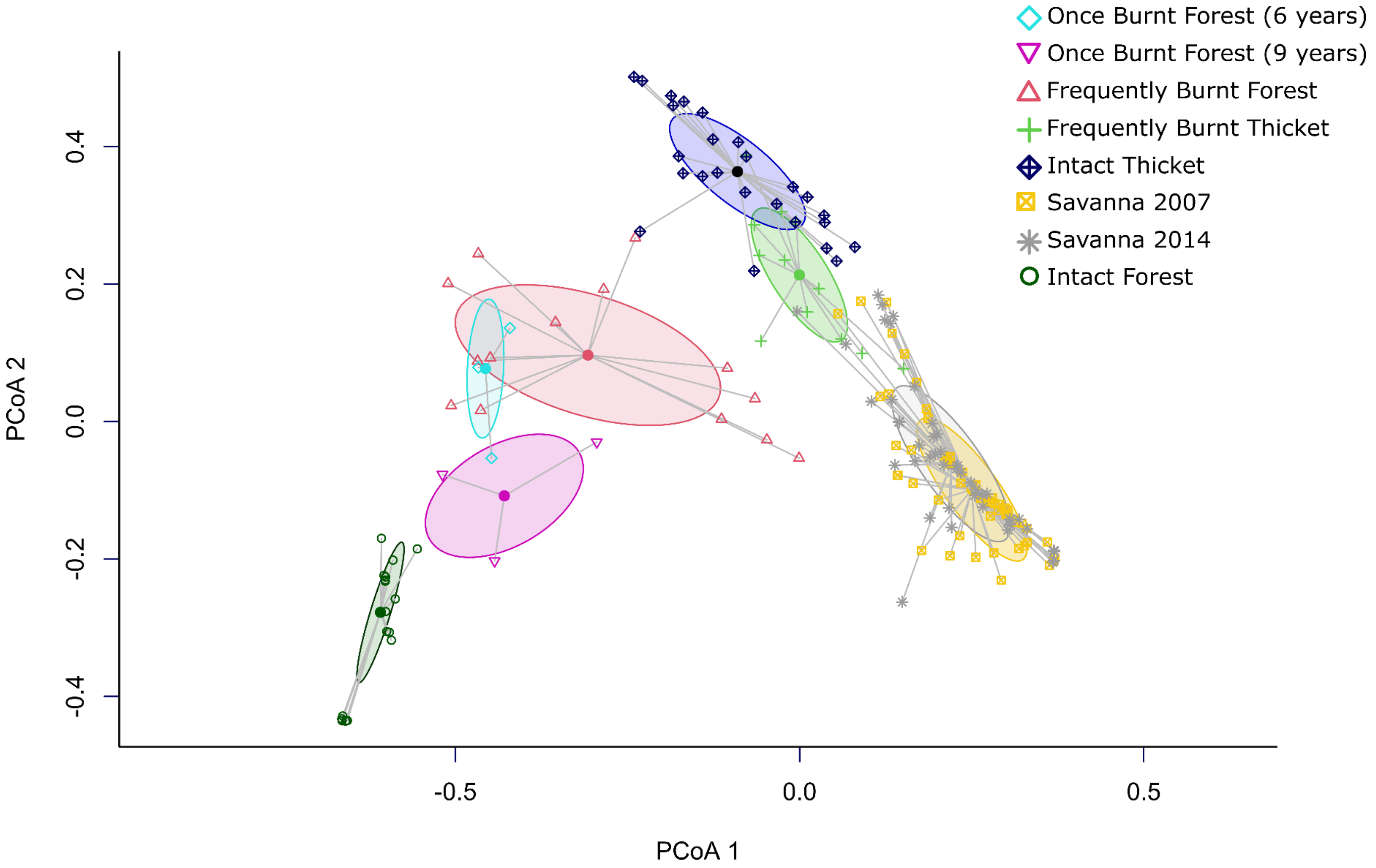


**Appendix Figure S3.** Principal Coordinates Analysis plot of the overall species type abundance in different vegetation types in Hluhluwe iMfolozi Park. Significant differences between groups for PERMANOVA and Pairwise PERMANOVA are shown in Appendix Table S5.
